# Temporal boundary gating of auditory sensitivity

**DOI:** 10.64898/2025.12.23.696169

**Authors:** Yuying Zhai, Haoxuan Xu, Hangting Ye, Peirun Song, Guifeng Zhai, Xuehui Bao, Ishrat mehmood, Nayaab Shahir Pandit, Yanyan Wang, Lingling Zhang, Pei Chen, Wanshun Wen, Gang Chen, Xuan Zhao, Yi Zhou, Xiongjie Yu

**Author notes:** Corresponding authors (X.Y.), (Y.Z.). These authors contributed equally (Yuying Zhai, Haoxuan Xu, Hangting Ye). Lead contact: Xiongjie Yu. **Author contributions:** Conceptualization, X.Y.; Methodology, X.Y.; Investigation and Data Curation, X.Y., Yuying Zhai, H.X., H.Y., P.S., G.Z., X.B., I.M., N.S.P., Y.W., L.Z., P.C., W.W., G.C., X.Z. and Yi Zhou; Writing – Original Draft, X.Y., Yuying Zhai, H.X. and H.Y.; Writing – Review & Editing, X.Y., Yuying Zhai, H.X., H.Y., P.S., G.Z., X.B., I.M., N.S.P., Y.W., L.Z., P.C., W.W., G.C., X.Z. and Yi Zhou; Funding Acquisition, X.Y., Yuying Zhai and Yi Zhou; Resources, X.Y.; Software, Yuying Zhai, H.X. and H.Y.; Validation and Visualization, Yuying Zhai, H.X. and H.Y.; Supervision, X.Y. **Declaration of interests:** The authors declare no competing interests.

## Abstract

Perception unfolds over time, but whether continuous sounds are sampled with uniform sensitivity is unknown. We combine human psychophysics, EEG and rodent neurophysiology to show that local auditory change detection is strongly gated by stimulus boundaries. In humans, brief perturbations inserted at different temporal positions within 0.5–1-s tones revealed an inverted U-shaped sensitivity profile: detection was attenuated near sound onset and offset and maximal mid-epoch, with EEG change responses showing a closely matching dependence on change timing. Electrocorticography in awake rats exhibited homologous temporal weighting, and analogous profiles for amplitude changes and visual motion demonstrated cross-feature and cross-modal generality. To uncover circuit mechanisms, we recorded single units along the inferior colliculus–medial geniculate body–auditory cortex pathway together with laminar local field potentials in A1. Onset-locked suppression of change responses emerged in midbrain and was progressively amplified in thalamus and cortex, whereas the full start–end profile was expressed in granular-layer alpha–gamma power. A simple biophysically grounded model in which onset responses saturate and cortical populations integrate over finite temporal windows recapitulates this pattern, explaining how stimulus boundaries disrupt prospective and retrospective integration and thereby degrade change detection near the beginning and end of sounds.

## Introduction

Our sensory experience unfolds in continuous streams, but our perception processes these streams in discrete temporal windows, ranging from tens to hundreds of milliseconds (Ulanovsky, Las et al. 2004, Murray, Bernacchia et al. 2014, Ding, Melloni et al. 2016, Ye, Song et al. 2025). This raises a fundamental question: is evidence from a continuous stimulus processed uniformly, or do different temporal segments contribute unequally to perception? In other words, does the brain treat a sustained event as a homogeneous whole, or does it prioritize certain time periods during processing (Rauschecker and Scott 2009, Nobre and van Ede 2018, Levi and Huk 2020, Horrocks, Rodrigues et al. 2024)? Most theoretical and computational accounts of temporal integration implicitly assume that, beyond well-known onset transients, sensitivity is approximately uniform over an event (Bogacz, Brown et al. 2006). Here, we challenge this assumption by investigating whether sensitivity to brief local changes is consistent across a sound’s duration or whether it varies systematically with temporal position.

A growing body of behavioral evidence suggests that sensory processing is temporally non-uniform. For example, when making judgments about dynamic intensity, observers integrate evidence with position-dependent weights, rather than uniformly over time (Nelken and Ulanovsky 2007, Oberfeld and Plank 2011, Baumgartner, Reed et al. 2017, Oberfeld, Fischenich et al. 2024). In the auditory system, onsets are particularly salient, capturing attention and disproportionately influencing perceptual reports compared to later segments (Naatanen and Winkler 1999, Naatanen, Paavilainen et al. 2007). This onset bias affects behaviors like speech perception, music listening, and environmental sound recognition (Rosen 1992, Schulze and Koelsch 2012, Zatorre and Baum 2012, Siedenburg 2019), suggesting that temporal position is a key factor in sensory coding. Additionally, theories of temporal attention propose that attention and expectation vary across an event’s unfolding, with different precision levels applied at different times (Large and Jones 1999, Matthews and Meck 2016, de Lange, Heilbron et al. 2018, Nobre and van Ede 2018). Onsets and offsets are recognized as event boundaries that segment continuous input into discrete perceptual episodes (Bregman 1990, Zacks, Speer et al. 2007). These findings imply that the brain may redistribute sensitivity over time, but the precise dynamics of this process across a continuous stimulus remain unclear.

At the neural level, auditory pathways exhibit pronounced response asymmetries across a stimulus (Song, Xu et al. 2025). Neurons typically fire strongly at sound onset and more weakly during sustained stimulation (Phillips, Hall et al. 2002, Hromadka, Deweese et al. 2008, Yu, Xu et al. 2009, Xu, Zhai et al. 2017, Rui, He et al. 2018, Zhai, Sun et al. 2019, Song, Zhai et al. 2021). Similarly, specialized coding mechanisms at sound termination have been identified in both subcortical and cortical circuits (Recanzone and Cohen 2010, Rauschecker 2015, Kopp-Scheinpflug, Sinclair et al. 2018, Chong, Anandakumar et al. 2020, Solyga and Barkat 2021, Song, Xu et al. 2024). Moreover, stimulus-specific adaptation and deviance detection reveal that recent stimulation strongly shapes responses to rare changes (Ulanovsky, Las et al. 2003, Nelken and Ulanovsky 2007, Antunes and Malmierca 2011, Taaseh, Yaron et al. 2011, Pérez-González, Hernández et al. 2012, Yaron, Hershenhoren et al. 2012, Xu, Yu et al. 2014, Song, Zhai et al. 2023, Du, Song et al. 2024, Gong, Song et al. 2024). These phenomena are framed within predictive coding and Bayesian models, in which the brain adjusts the precision of predictions over time (Friston 2005, Feldman and Friston 2010, de Lange, Heilbron et al. 2018, Gifford, Sperling et al. 2019, Du, Xu et al. 2025). However, how these neural asymmetries translate into position-dependent sensitivity to local changes is still unknown. We aim to clarify this by characterizing whether continuous sounds are processed uniformly or follow a structured temporal weighting profile tied to event boundaries.

To address this gap, we used a local change-detection paradigm where brief perturbations are embedded at different positions within continuous auditory stimuli. If processing were temporally uniform, detection performance and neural responses would remain constant across perturbation timing. Any systematic deviations reveal how sensitivity is redistributed across the stimulus. By combining human psychophysics and EEG with rat electrocorticography (ECoG), single-unit recordings along the inferior colliculus–medial geniculate body–auditory cortex (IC–MGB–AC) pathway, and laminar local field potentials (LFPs) in A1, we map how sensitivity to local changes varies over time across the auditory hierarchy. Behaviorally, we find that local change detection is strongly suppressed near sound onset and offset but peaks in the middle of the stimulus—a robust “start–end effect” that generalizes across acoustic features and is mirrored in a visual task. Neurally, onset-linked suppression emerges in midbrain and thalamus and is progressively amplified in cortex, whereas offset-linked suppression is weak and variable at the single-unit level but is prominently expressed in population field potentials and layer-specific oscillations. These convergent results identify temporal boundary gating as a fundamental, evolutionarily conserved organizing principle: the brain down-weights sensory evidence near event boundaries while prioritizing the more stable mid-epoch. This challenges current theories of temporal integration (Hasson, Yang et al. 2008, Murray, Bernacchia et al. 2014, DiTullio, Parthiban et al. 2023), predictive coding (Chaudhuri, Knoblauch et al. 2015), and hierarchical timescales (Horrocks, Rodrigues et al. 2024) by proposing a non-uniform temporal weighting scheme that redistributes sensitivity across the stimulus, particularly near event boundaries.

## Results

We investigated whether sensitivity to brief local changes in a continuous stimulus is uniform across time or shaped by stimulus boundaries. To address this, we combined human psychophysics and EEG with rat ECoG, Neuropixels recordings along the IC–MGB–AC pathway, and laminar LFPs in A1.

### Start–End Effect in Human Perception

We investigated whether continuous sounds are processed uniformly or if different temporal segments contribute unequally to perception. In a local change-detection task (Fig. 1A), human participants listened to 1-s, 1-kHz pure tones with a ∼20-ms frequency perturbation (Δf) inserted at one of 11 temporal positions (50–950 ms) or omitted (control). Detection performance showed a clear inverted U-shaped profile, with sensitivity lowest at sound onset (50–100 ms) and offset (850–950 ms) and maximal sensitivity in the middle of the stimulus (∼250–500 ms). Sensitivity at mid-stimulus positions was significantly higher than at both early and late positions (p < 0.01, FDR-corrected), demonstrating temporal non-uniformity in change detection (Fig. 1B).

**Figure 1.**
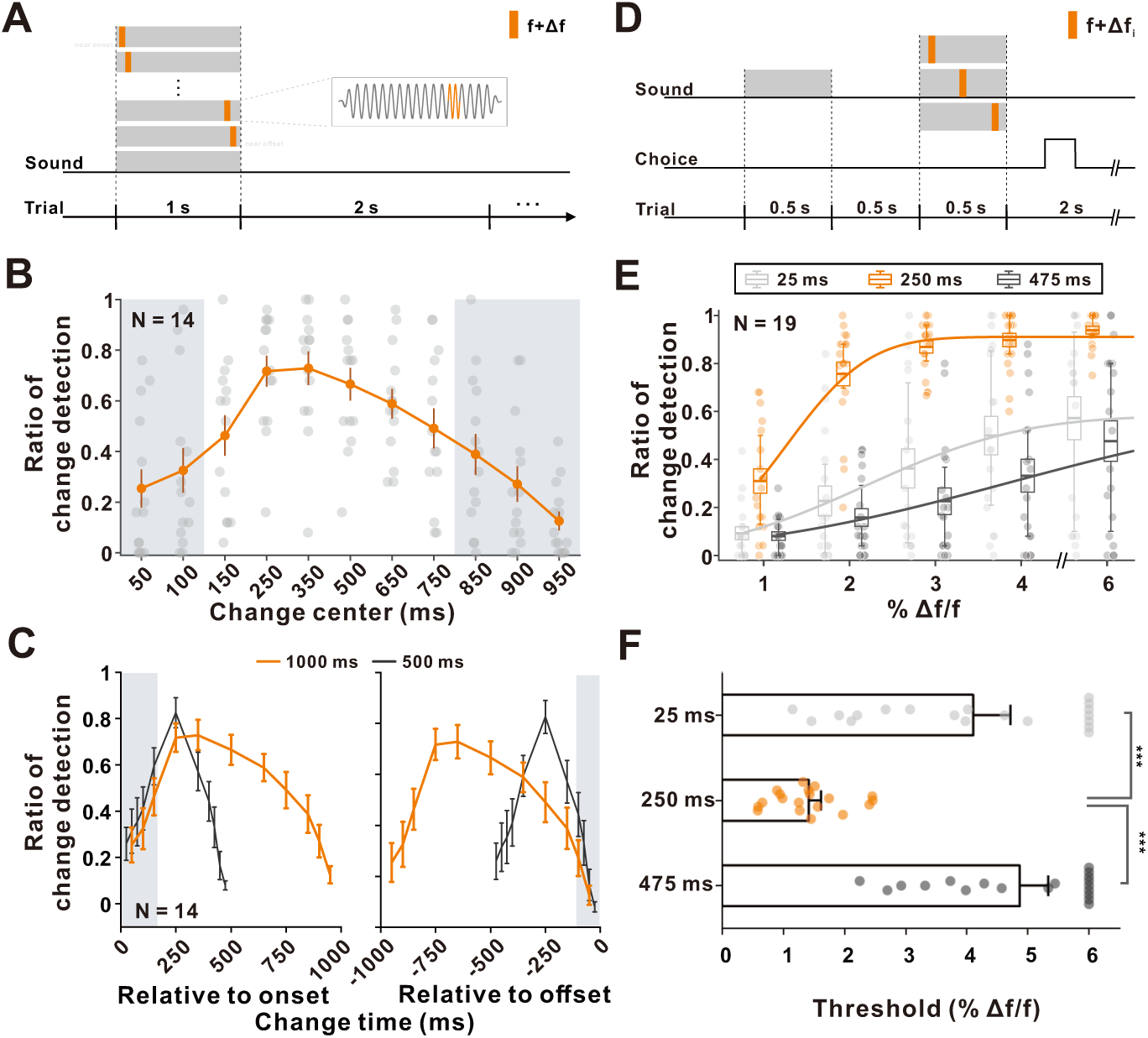
Psychological and behavioral performance in the local change-detection task. **(A)** Schematic of the local change-detection paradigm. Each trial contained a continuous 1-s, 1-kHz tone (gray), followed by 2 s of silence. A brief frequency perturbation (orange; f + Δf) was embedded at one of several predetermined temporal positions during the 1-s tone. Participants responded via left or right keypresses to indicate whether a change occurred on each trial. The right inset provides an example illustration of the local frequency increase (schematic only). **(B)** Change-detection performance across temporal positions for the 1-s tone condition (N = 14). Each point represents an individual participant; boxplots show the mean (central line), ±1 SEM (box edges), and 25th–75th percentiles (whiskers). Shaded gray mark temporal positions where detection performance was significantly reduced relative to the position with maximal mean performance (one-way ANOVA across positions followed by post-hoc comparisons, p < 0.05). **(C)** Temporal sensitivity profiles for 500-ms (black) and 1000-ms (orange) tones, plotted relative to sound onset (left panel) or offset (right panel). Shaded gray regions indicate time ranges where detection performance did not differ significantly across adjacent positions (N = 14; mean ± SEM). **(D)** Delayed match-to-sample task. Participants compared two 500-ms tones (1 kHz) separated by a 500-ms silent interval. The second tone contained a 20-cycle frequency increase at one of three positions (25 ms, light gray; 250 ms, orange; 475 ms, dark gray), with Δf magnitude varied (0%, 1%, 2%, 3%, 4%, 6%) across trials. Participants judged whether the two tones matched (Psychological Experiment, Frequency Session 1). **(E)** Change-detection performance for perturbations at 25 ms (light gray), 250 ms (orange), and 475 ms (dark gray). Transparent dots indicate individual participant data (N = 19); boxplots summarize the mean (central line), ±1 SEM (box edges), and 25th–75th percentiles (whiskers). A Gaussian function was fit to the pooled data to visualize the psychometric relationship. **(F)** Distribution of individual detection thresholds, defined as the Δf producing 60% detection probability, for the three perturbation positions. Each dot represents one participant; bars indicate mean ± SEM. Significant differences between conditions are marked with asterisks (***, p < 0.001).

Next, we tested whether this effect is anchored to sound onset or offset. We compared detection across two stimulus durations (500 ms and 1000 ms) and realigned performance either to onset or offset. We found that sensitivity reductions at both early and late positions were tied to stimulus boundaries, not to absolute time, confirming that both the early and late components of the start–end effect are anchored to stimulus boundaries (Fig. 1C).

To more precisely measure sensitivity, we used a delayed match-to-sample paradigm. Participants compared two 500-ms tones, one with a 20-ms frequency shift at the start, middle, or end (Fig. 1D). Sensitivity was highest in the middle of the tone (mean 1.42% Δf/f) and significantly lower at both the start (4.10%) and end (4.87%) (Fig. 1E–F), confirming that local change detection is most sensitive mid-epoch and attenuated near stimulus boundaries.

To assess the generalizability of this effect across different sensory features and modalities, we replicated the inverted U-shape with amplitude perturbations (Supplementary Fig. 1) and in a visual task where participants detected a luminance increase in a moving bar (Supplementary Fig. 2). These results show that temporal non-uniformity extends beyond auditory stimuli, influencing visual processing as well and supporting the idea of cross-modal temporal weighting.

### EEG Signatures of Temporal Boundary Gating in Humans

We next investigated whether the behavioral start–end effect is reflected in macroscopic neural activity. In a passive-listening EEG experiment, 30 participants listened to 1-s, 1-kHz tones, each containing a ∼20-ms, 8% frequency increase at one of 11 temporal positions (50–950 ms) or no change (control). Participants were not required to respond, allowing us to isolate stimulus-locked change responses. Event-related potentials (ERPs) showed clear position-dependent change responses. At the T8 electrode, perturbations occurring mid-stimulus (e.g., 200 ms) evoked large deflections relative to control, whereas early (50 ms) and late (950 ms) perturbations elicited significantly smaller responses (Fig. 2A). A cluster-based permutation test on global field power (GFP) identified a 60–220 ms window after perturbation onset as the change-response period (Methods). Within this window, we quantified response magnitude using a root-mean-square (RMS) metric and defined ΔRM as the difference between change and control conditions for each temporal position.

**Figure 2.**
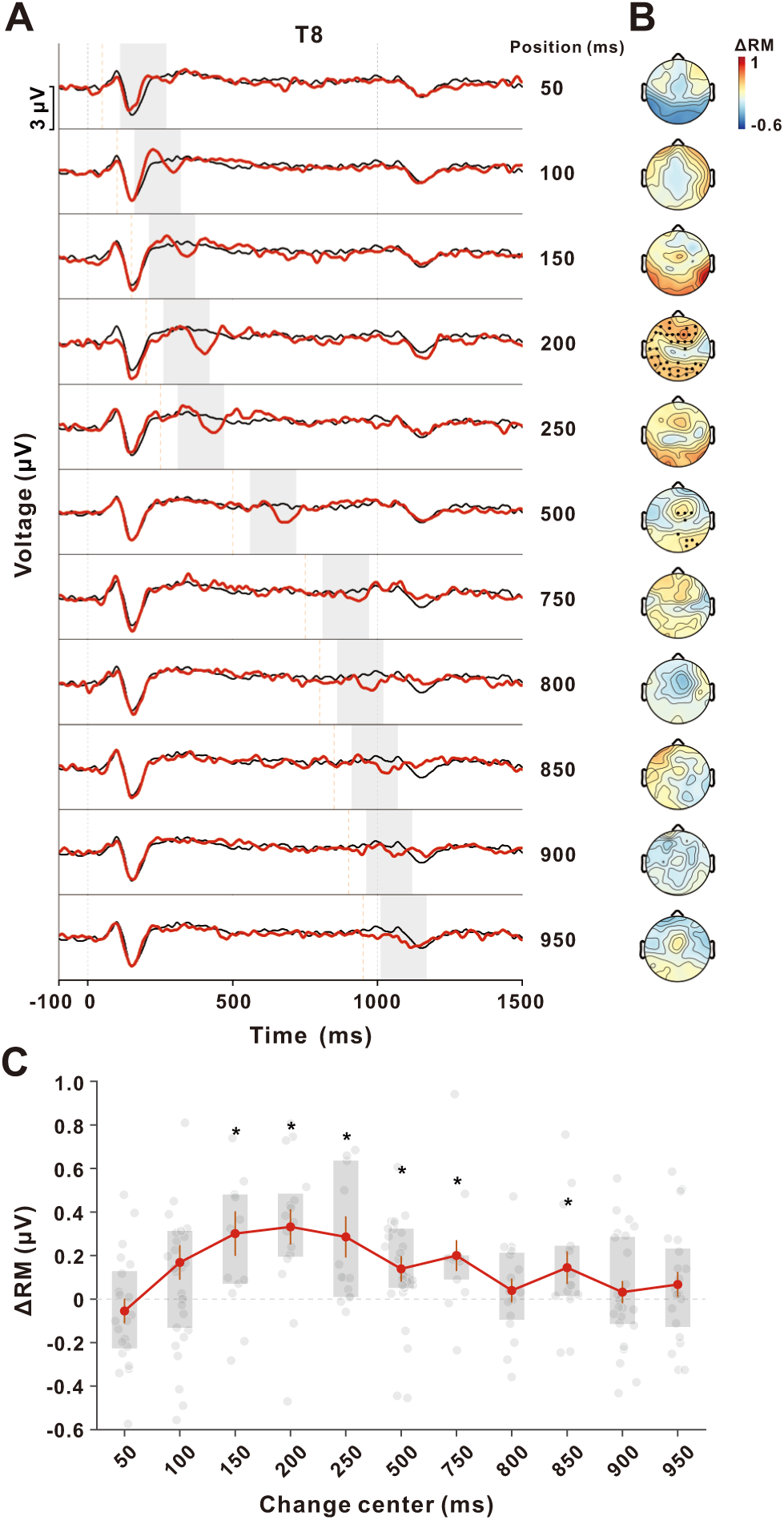
EEG change responses during the change-detection task. **(A)** Grand-averaged EEG responses recorded at electrode T8 in the Human EEG Experiment (N = 30). Red color traces show responses to each perturbation condition, and black traces show the no-change control. Light orange dashed lines mark the times of the perturbations (50–950 ms). Light gray dashed lines denote sound onset and offset. The shaded region (60–220 ms after perturbation onset) indicates the time window used to compute the response magnitude (RM), defined based on significant cluster difference between the control condition and 50% change position (see Methods, p < 0.05, permutation test). **(B)** Scalp topographies of ΔRM (response magnitude difference relative to control) for each perturbation position from top to bottom (50, 100, 150, 200, 250, 500, 750, 800, 850, 900, or 950 ms). ΔRM was computed as RM_change_ − RM_control_. Black dots mark electrodes showing significant change responses compared with control (one-tailed paired t-test, FDR-corrected, p < 0.05). **(C)** Group-level ΔRM across the 11 perturbation positions. For each participant, ΔRM was computed for each position and then averaged across EEG channels. Gray dots represent individual participants (N = 30). Red solid line with filled circles denote the mean ΔRM across participants at each position, with vertical error bars indicating ±1 SEM. Shaded rectangles indicate the interquartile range (IQR, 25th–75th percentiles). Asterisks indicate positions where ΔRM was significantly greater than zero (one-tailed paired t-test, p < 0.05). The light-gray dashed line denotes ΔRM = 0.

Topographic maps of ΔRM (Fig. 2B) revealed widespread scalp activity for mid-stimulus perturbations, with maximal responses over frontotemporal electrodes and weaker responses for early and late perturbations. When ΔRM was averaged across channels, the resulting profile was inverted U-shaped (Fig. 2C): population-level change responses peaked for mid-stimulus positions and declined toward both onset and offset. ΔRM values at early (50 ms) and late (950 ms) positions were significantly smaller than at mid-stimulus positions (p < 0.05, one-tailed paired t-tests; Fig. 2C). These results provide a clear neural correlate of temporal boundary gating in human EEG, with scalp-recorded change responses exhibiting the same start–end effect observed behaviorally.

### Cross-Species Evidence for the Start–End Effect in Rats

To determine whether temporal boundary gating is conserved across species, we recorded electrocorticography (ECoG) from the auditory cortex of awake, head-fixed rats (see setup in Fig. 3A). Rats passively listened to a paradigm based on the same logic as the human EEG task, where a brief local perturbation was embedded within a continuous sound. The stimuli consisted of 1-s complex tones (4, 8, 16, and 32 kHz components), with a ∼4-ms frequency increase (∼3.6%) inserted at one of 13 temporal positions (25–975 ms) or omitted (control).

**Figure 3.**
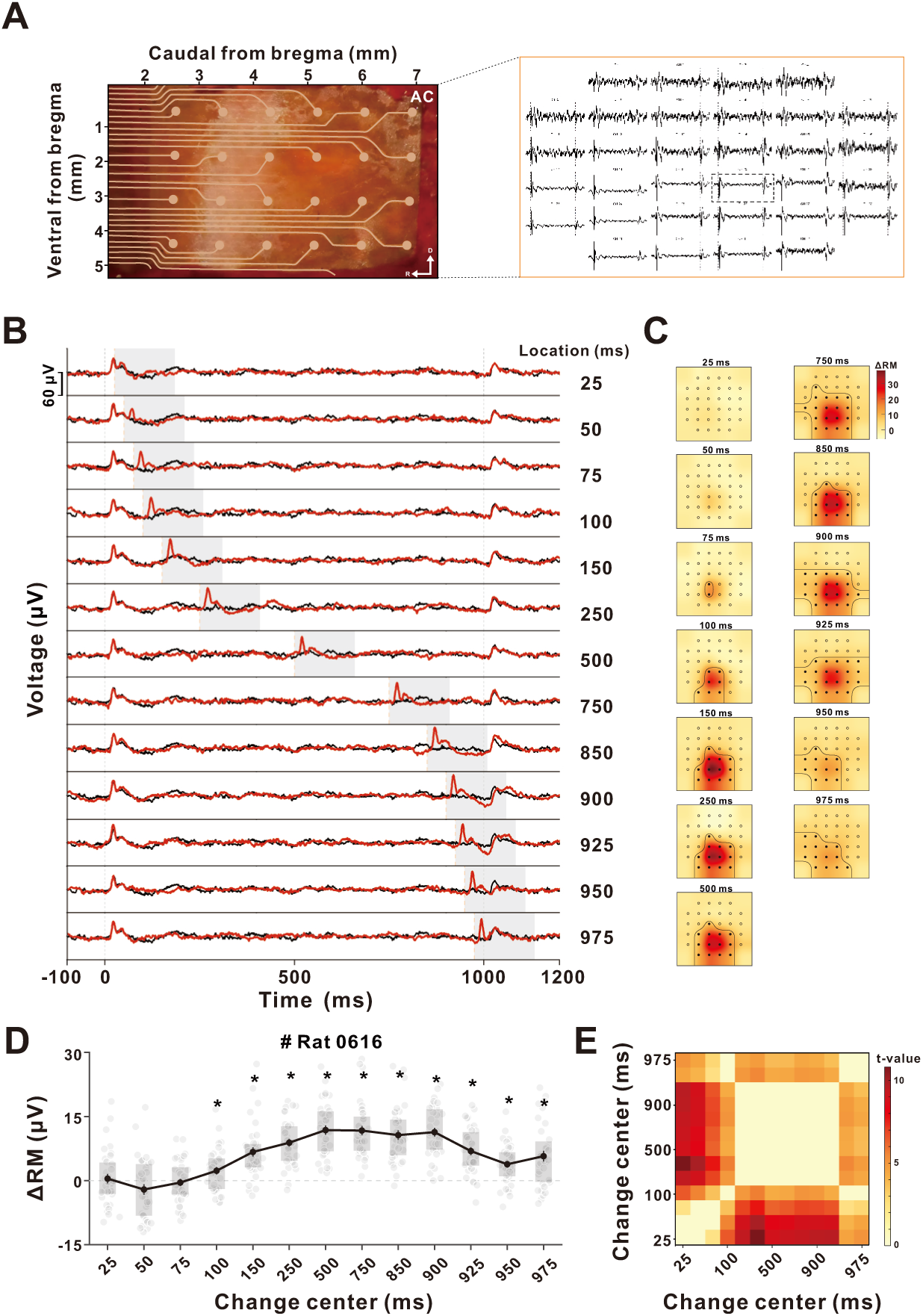
ECoG change responses during the local change-detection task. **(A)** Left: Photograph of the ECoG array implanted over the auditory cortex (AC) in an example rat. Right: Raw ECoG traces recorded from the full grid during stimulus presentation. The dashed box highlights channel #27, shown in (B). **(B)** Example ECoG responses from channel #27 of Rat 0616 during the change detection task in Rat ECoG Experiment, Session 1. Black traces show responses to the no-change control condition, and red color traces correspond to different perturbation positions. Orange dashed vertical lines indicate the onset times of perturbations (25–975 ms) and light gray dashed lines indicate the onset and offset of sound stimuli. The shaded region (0–160 ms after perturbation onset) indicates the time window used to compute RM value, based on significant differences observed between the control and 50% change conditions (p < 0.05, permutation test). **(C)** Topographical distribution of ΔRM across the ECoG array for each perturbation position. ΔRM was computed as RM_change_ - RM_control_. Electrode sites enclosed by black contours show significant change responses (one-tailed paired t-test, FDR-corrected, p < 0.05). **(D)** ΔRM across perturbation positions for Rat 0616, averaged over the 16 channels with the largest onset responses (see Methods). Gray dots represent ΔRM values from individual trials. Black solid line with filled circles denote the mean ΔRM at each perturbation position, with vertical error bars indicating ±SEM. Shaded rectangles indicate the IQR with 25th–75th percentiles. The light gray dashed line denotes ΔRM = 0. Asterisks indicate positions where ΔRM was significantly greater than zero (one-tailed t-test, p < 0.05). **(E)** Pairwise t-value matrix comparing ΔRM across all perturbation positions. Each matrix cell shows the t-value for the comparison between two positions; nonsignificant comparisons (p ≥ 0.05) are masked. Darker colors indicate larger t-values.

In a representative channel, perturbations near the onset (25–50 ms) evoked minimal change-related activity compared to control, while mid-stimulus perturbations elicited clear response deflections. Offset-proximal perturbations (950–975 ms) also triggered responses, though these were weaker than those at mid-sequence positions (Fig. 3B). ΔRM, calculated within a 0–160 ms post-change window, showed strong responses centered around mid-sequence positions (150–925 ms) and reduced responses near both onset and offset (Fig. 3C). When averaged across the 16 channels with the largest onset responses, ΔRM followed an inverted U-shaped profile: responses rose sharply from early positions, remained elevated through the central portion of the stimulus, and dropped off sharply near the end (Fig. 3D). For most mid-sequence positions, ΔRM was significantly greater than zero (p < 0.05, one-tailed t-tests), while early and late positions exhibited reduced ΔRM values.

Pairwise t-tests on ΔRM across change times revealed a characteristic boundary-centered structure (Fig. 3E). Comparisons including onset-proximal (25–100 ms) or offset-proximal (950–975 ms) changes survived significance masking, yielding large t-values. In contrast, pairwise comparisons within the mid-sequence positions (150–925 ms) were largely non-significant, forming a central region with no surviving t-values. This pattern indicates that boundary-proximal changes elicit distinct responses compared to mid-epoch changes, while mid-epoch responses are more homogeneous. Two additional rats showed similar profiles (Supplementary Fig. 3), and analogous start–end effects were observed when the perturbation was implemented as an amplitude increase instead of a frequency shift (Supplementary Fig. 4). Thus, the start–end effect is a robust, cross-species property of auditory cortical population activity.

### Emergence of the Start–End Effect Along the Auditory Pathway

We sought to identify where along the ascending auditory pathway temporal boundary gating first emerges and how it is transformed across stages. Using Neuropixels probes, we recorded extracellular activity in the inferior colliculus (IC), medial geniculate body (MGB), and primary auditory cortex (AC) of awake rats during the same complex-tone local-change paradigm used for ECoG (Fig. 3).

To illustrate how temporal boundary gating manifests at the level of individual neurons, we first examined single-neuron responses in AC (Fig. 4). In one example neuron, perturbations near the onset produced little additional spiking relative to control, whereas perturbations from ∼150 ms onward elicited clear increases in firing (Fig. 4A). Δ firing rate (ΔFR; change minus control, computed in a 0–100 ms post-perturbation window) rose steadily from early positions and remained high across the mid-to-late portion of the sequence, with a visually apparent plateau spanning ∼750–950 ms (Fig. 4C). The very latest perturbations (950–975 ms) produced a modest decline relative to this high-response interval. Pairwise tests confirmed significantly reduced responses at early positions and the relative homogeneity of firing across the plateau region (Fig. 4D).

**Figure 4.**
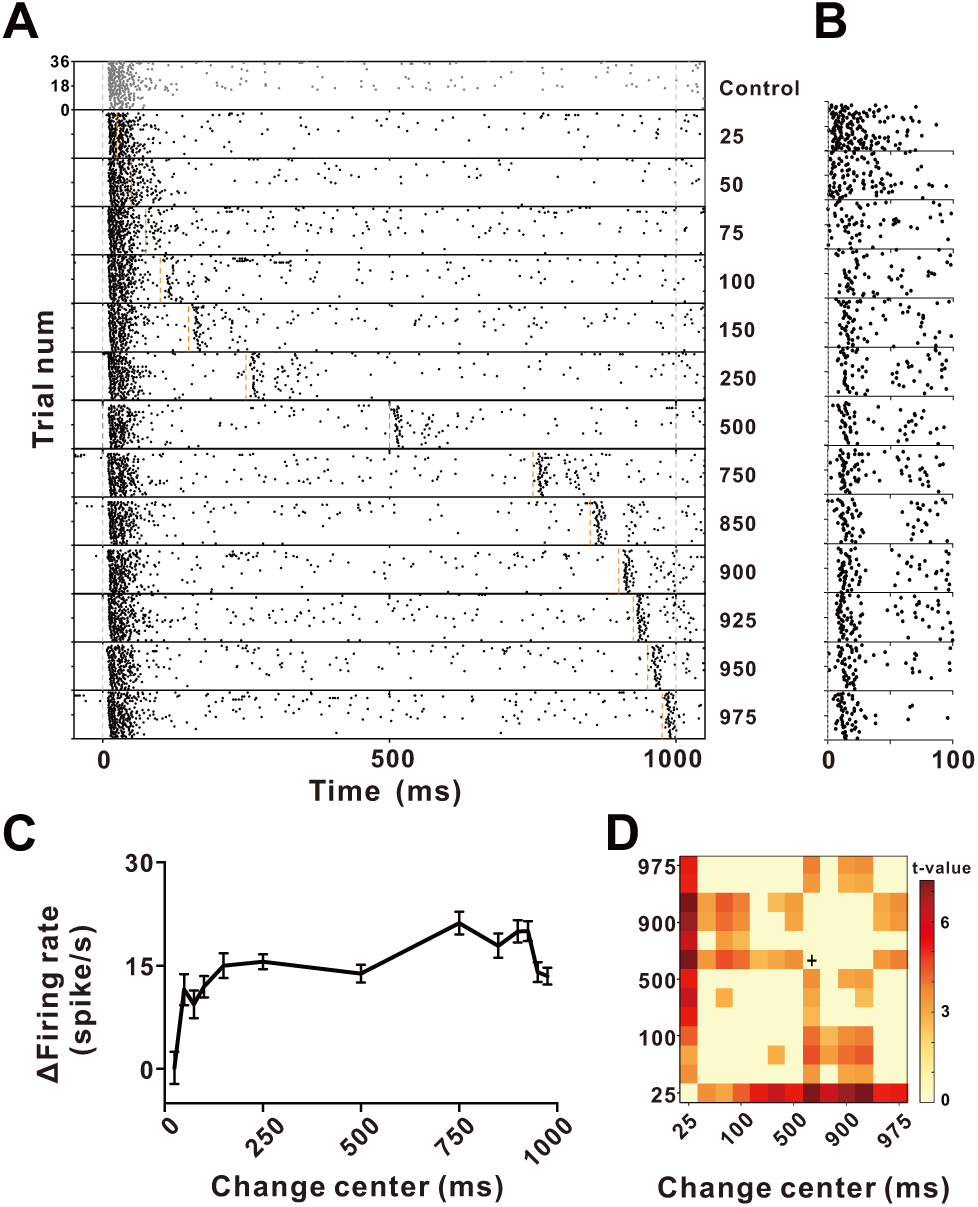
Example neuronal responses in auditory cortex (AC) to local changes. **(A)** Raster plot of spike responses from an example AC neuron during the change-detection task (Rat Extracellular Experiment). Orange dashed lines mark the tested perturbation times (25–975 ms). Gray dashed lines mark sound onset and offset. **(B)** Responses from the same neuron, realigned to each perturbation time (0–100 ms window). **(C)** Δ firing rate (ΔFR; change minus control, computed in a 0–100-ms window after perturbation onset) as a function of perturbation time. ΔFR rose sharply from early positions, reached a broad mid-sequence maximum, and then declined modestly at the latest positions—showing weaker end-related suppression compared with the strong reduction near stimulus onset. Error bars indicate ±SEM. **(D)** Pairwise t-value matrix comparing ΔFR across perturbation positions. Each cell reflects the t-value for the comparison between two positions; non-significant values (p ≥ 0.05) are masked. The cross symbol marks the position of the maximum ΔFR. The pattern confirms significantly reduced responses at early positions and comparatively smaller reductions at the latest positions.

In IC, many neurons responded to local changes primarily with suppression rather than excitation (Supplementary Fig. 5), with ΔFR decreasing at later positions or showing no significant modulation. To focus population analyses on units that contributed positively to change encoding, we restricted analyses to neurons whose maximal ΔFR was significantly greater than zero.

Across this subset, we characterized each neuron’s temporal “plateau” of maximal responsiveness. Neurons were sorted by the position of their maximal ΔFR, and for each neuron, we identified the left (green line) and right (blue line) boundaries of the plateau—the earliest and latest positions whose responses were statistically indistinguishable from the maximum (Fig. 5A–C). Positions earlier than the green line or later than the blue line showed significantly reduced ΔFR, defining start- and end-related suppression relative to the neuron’s preferred temporal window. This analysis revealed a clear hierarchical trend. In AC, most neurons showed strong onset-locked suppression (Fig. 5A): ΔFR was significantly reduced for early positions, then rose to a sustained plateau from mid-sequence onward, with only a minority of neurons exhibiting robust suppression near offset. MGB neurons showed a similar but weaker pattern (Fig. 5B), whereas IC neurons displayed the least pronounced onset suppression and relatively flat profiles across later positions (Fig. 5C).

**Figure 5.**
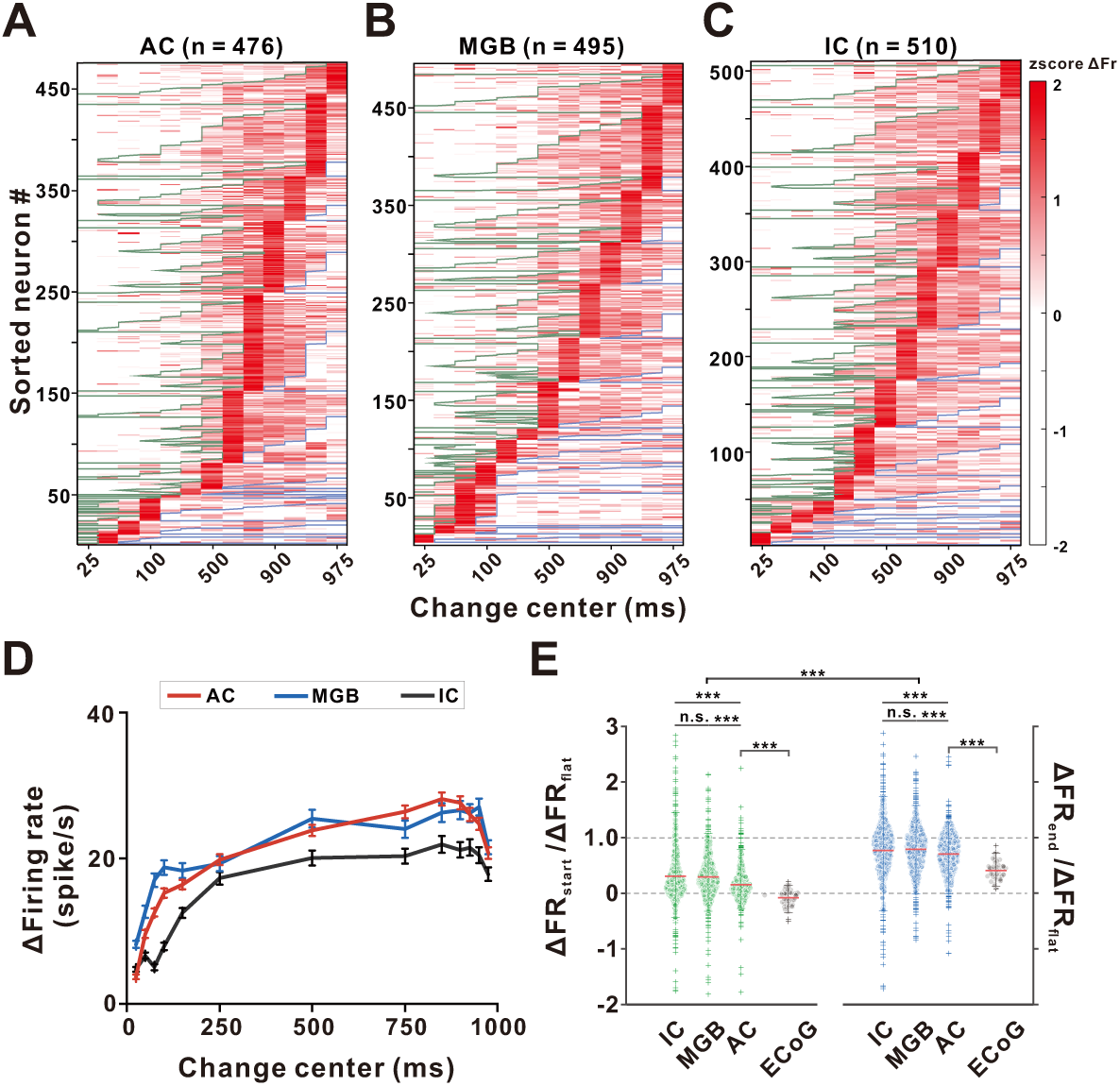
Population neuronal responses in AC, MGB and IC during local change detection. **(A-C)** Heatmaps show z-scored Δ firing rates (ΔFR; change minus control) for neurons in the AC (n = 476), MGB (n = 495), and IC (n = 510), sorted by the temporal position of each neuron’s maximal response. Each row corresponds to a single neuron, and each column to one of the 13 perturbation times. Warmer colors indicate stronger ΔFR. Overlaid green and blue boundaries indicate the temporal limits within which each neuron’s firing remains statistically indistinguishable from its maximum response. Specifically, the green line marks the earliest perturbation time whose ΔFR does *not* differ significantly from the maximum (i.e., all earlier positions show significantly lower responses), whereas the blue line marks the latest perturbation time still statistically indistinguishable from the maximum (i.e., all later positions show significantly lower responses). Thus, the interval between these two boundaries defines the neuron’s plateau period, during which firing is maintained at levels not significantly different from the maximal ΔFR (see Methods for details). **(D)** Average ΔFR across perturbation positions for neurons in AC (red), MGB (blue), and IC (black). Error bars: ±SEM. **(E)** Ratios of Δ firing rates (ΔFR) at boundary-proximal positions relative to each neuron’s plateau response. For spike data (green and blue), ΔFR at 25 ms (start) and 975 ms (end) were divided by the neuron’s mean ΔFR (“flat”) during its plateau period, defined individually for each neuron by the green and blue boundaries in panels A–C. For ECoG data (gray), ΔRM values at the same two positions (25ms and 975 ms) were normalized by the maximum ΔRM (“flat”) across all perturbation times for that channel. Each transparent dot represents one unit (spikes) or one channel (ECoG); red horizontal lines denote group means. Gray dashed lines indicate ratios of 0 and 1. Statistical comparisons across groups and conditions are shown (***p < 0.001; n.s., not significant).

Population tuning curves confirmed these regional differences (Fig. 5D). In AC, mean ΔFR increased with later change positions and saturated around ∼750 ms; in MGB, the rise was more gradual and plateaued slightly earlier; in IC, ΔFR increased modestly and remained at lower overall magnitude. To quantify boundary effects relative to each neuron’s plateau, we computed the ratios ΔFR_25ms_/ΔFR_flat_ (start) and ΔFR_975ms_/ΔFR_flat_ (end), where ΔFR_flat_ is the mean ΔFR over the neuron’s plateau positions (Fig. 5E). Across regions, start ratios were significantly smaller than end ratios (p < 0.001, ANOVA with post-hoc tests), indicating stronger onset than offset suppression. Moreover, start ratios decreased systematically along the hierarchy—from IC to MGB to AC—demonstrating progressive amplification of onset-related suppression in cortex. End ratios were closer to the plateau value and showed only modest differences across regions. Importantly, ECoG end ratios were substantially lower than those observed in single units, suggesting a more pronounced population-level reduction at stimulus offset (Fig. 5E). These results suggest that the start effect—a transient reduction of sensitivity following sound onset—emerges in midbrain and thalamus and is progressively strengthened in cortex, whereas the end effect is weaker and more heterogeneous at the single-unit level.

### Cortical Population and Laminar Organization of the Start–End Effect

The ECoG data demonstrated a robust start–end effect at the cortical population level, whereas single-unit analyses revealed a dominant start effect with comparatively weaker and more variable end effects. To reconcile these findings, we analyzed laminar local field potentials (LFPs) extracted from the same Neuropixels recordings in A1, using depth-resolved channels to assess layer-specific population activity while presenting the same local-change stimuli (Fig. 6).

**Figure 6.**
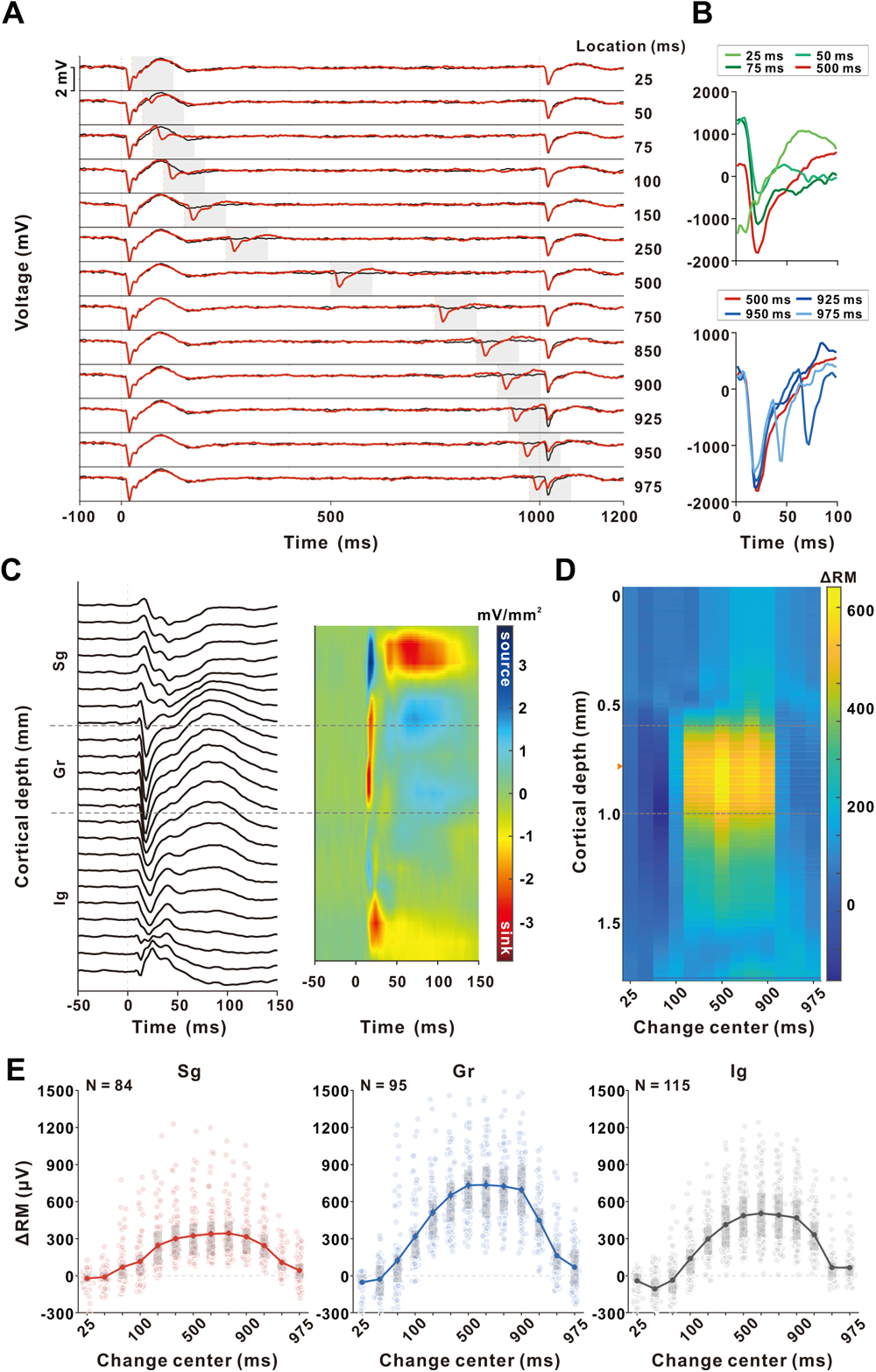
Laminar LFP signatures of temporal boundary gating in A1. **(A)** Example LFP responses from a single channel in A1 during the local change-detection paradigm. Black traces show responses in the control condition; red traces show responses when a local change occurred at one of 13 temporal positions (25–975 ms). Gray vertical dashed lines indicate sound onset and offset, and orange vertical dashed lines mark the timing of the local change. Gray shaded regions indicate the 0–100 ms post-change analysis window used to compute relative magnitude change (RM). **(B)** Perturbation-aligned LFP traces (0–100 ms) from the same penetration. Top: early-change responses (25–75 ms) grow progressively larger as the perturbation occurs farther from sound onset. Bottom: late-change responses (925–975 ms) weaken as the perturbation approaches sound offset. Together, these patterns mirror the start–end suppression observed in behavior and ECoG. **(C)** Current-source density (CSD) analysis used to assign laminar boundaries. Left: LFP responses to a brief noise burst recorded across probe depth. Right: CSD map derived using a five-point spatial derivative. Dashed horizontal lines mark supragranular (Sg), granular (Gr), and infragranular (Ig) layer boundaries, determined from the earliest current sink in the granular layer (Methods). **(D)** Depth-resolved ΔRM map for the same penetration shown in (C). For each channel, ΔRM was computed as the change-condition RM minus the control-condition RM. Warm colors indicate larger LFP responses to mid-sequence changes, whereas early and late changes produced weak ΔRM across layers. The orange triangle marks the channel shown as the example in panel (A). **(E)** Population laminar ΔRM across penetrations. For each penetration, channels were assigned to Sg, Gr, or Ig using the CSD boundaries in (C). Within each layer, ΔRM values were extracted every 0.1 mm and plotted for all penetrations (Sg: *N* = 84 channels; Gr: *N* = 95; Ig: *N* = 115). Gray dots represent individual channels. Solid lines with filled circles denote the mean ΔRM across channels at each change position, and vertical error bars indicate ±1 SEM. Shaded rectangles indicate the IQR with 25th–75th percentiles. Across layers, ΔRM was minimal for early and late changes and maximal for mid-sequence changes, with the granular layer exhibiting the largest modulation.

In an example penetration, LFPs from a single channel showed clear dependence on change timing: perturbations near onset (25–100 ms) or offset (925–975 ms) produced small deflections, whereas mid-stimulus changes evoked larger responses (Fig. 6A). When traces were realigned to perturbation onset, responses to early changes (25–75 ms) grew progressively larger as the changes occurred farther from sound onset (Fig. 6B, top). In contrast, responses to late changes (925–975 ms) weakened as the perturbation approached sound offset (Fig. 6B, bottom), paralleling the behavioral and ECoG start–end effect.

We assigned recording contacts to supragranular (Sg), granular (Gr), and infragranular (Ig) layers based on current-source density (CSD) profiles evoked by brief noise bursts (Fig. 6C). ΔRM maps plotted as a function of depth and change position revealed a pronounced inverted U-shaped profile in the granular layer: ΔRM was minimal for early and late changes and maximal for mid-sequence changes (Fig. 6D). Similar but weaker modulation was also observed in Sg and Ig. Population averages confirmed that all three layers exhibit the start–end effect, with the most pronounced modulation in Gr (Fig. 6E). In each layer, ΔRM rose from early positions, peaked for mid-stimulus changes, and declined again near offset. The dynamic range of this inverted U-shaped profile was greatest in Gr, intermediate in Ig, and smallest in Sg.

Thus, cortical population activity—particularly in the thalamorecipient granular layer—exhibits a strong non-uniform temporal weighting of local changes, even when single-unit firing shows limited end-related suppression. Consistent with this, depth-resolved LFPs in MGB and IC (Supplementary Fig. 6) exhibited robust start effects but little systematic modulation near offset, mirroring the spike data from these regions. Thus, the full start–end pattern emerges at the level of cortical population activity, most prominently in the A1 granular layer.

### Oscillatory Dynamics Underlying Temporal Boundary Gating

Finally, we examined how the start–end effect is expressed in the frequency domain. Power changes were quantified as ΔP/P_control_, representing the relative power increase compared with control trials. Time–frequency analysis of cortical LFPs revealed position-dependent modulations of band-limited power (Fig. 7). In an example penetration, spectrograms showed that power in the 8–64-Hz range was markedly reduced when changes occurred near onset (25–50 ms) or offset (925–975 ms), but was strongly enhanced for mid-stimulus changes (250–750 ms) across channels spanning the cortical laminae (Fig. 7A). After assigning channels to supragranular (Sg), granular (Gr), and infragranular (Ig) layers based on current-source density (CSD), we averaged time–frequency representations across penetrations. All three layers displayed the same qualitative profile: power in the low-gamma range (∼30–60 Hz), and to a lesser extent in 8–30 Hz, was strongly enhanced for mid-sequence perturbations compared to early or late ones, with the largest modulation in the granular layer (Fig. 7B).

**Figure 7.**
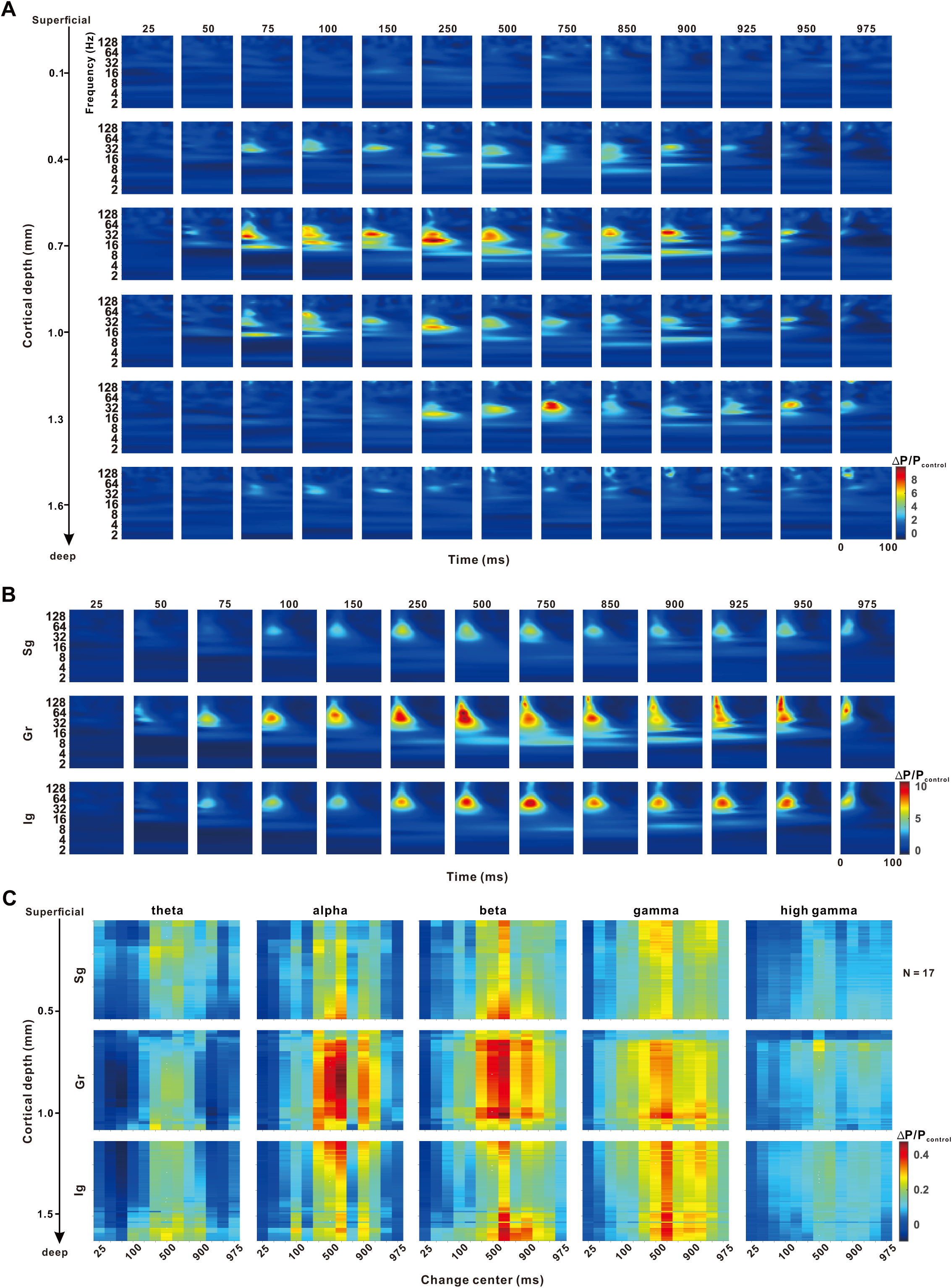
Laminar LFP time–frequency responses to local changes. (A) Example laminar time–frequency representation (TFR) from a single penetration. Spectral power changes are shown for each perturbation position (25–975 ms). Channels are ordered from superficial to deep cortex (approximate depths labeled on the left). For each channel and each change position, the power spectrum was computed and normalized to the control condition (ΔP/P_control_ = (P_change_ − P_control_) / P_control_). Each panel displays the 0–100 ms post-change window used for quantifying change-related power. (B) Across-penetration mean laminar TFRs. For each penetration, channels belonging to superficial (Sg), granular (Gr), and infragranular (Ig) layers were identified using depth estimates and CSD-based boundaries, and the median ΔP/P_control_ within each layer was computed. Heatmaps show the across-penetration average (N = 17), summarizing layer-specific change-related spectral responses across perturbation positions. (C) Depth-aligned laminar power across canonical frequency bands. For each penetration, ΔP/P_control_ was averaged within five frequency ranges (theta: 4–8 Hz; alpha: 8–14 Hz; beta: 14–30 Hz; gamma: 30–80 Hz; high-gamma: 80–150 Hz) during the 0–100-ms post-change window. Channels were depth-aligned across penetrations using CSD-defined borders, allowing direct comparison of Sg, Gr, and Ig layers despite variability in absolute depth. Heatmaps show the across-penetration average (N = 17), with the vertical axis indicating laminar position (superficial at top, deep at bottom).

To further characterize these effects across canonical frequency bands, we aligned channels across penetrations based on CSD-defined layer boundaries and computed depth-resolved band-limited power as a function of change position (Fig. 7C). Alpha, beta, and gamma bands in all layers exhibited clear inverted U-shaped profiles, with maximal power for mid-stimulus changes and reduced power near onset and offset. In contrast, theta and high-gamma bands showed much weaker position dependence.

These findings indicate that temporal boundary gating is implemented not only in mean LFP amplitude but also in coordinated oscillatory dynamics. Low- and mid-frequency oscillations (alpha–gamma) across cortical layers collectively emphasize mid-epoch perturbations while down-weighting those near stimulus boundaries, providing a population and spectral mechanism for the non-uniform temporal weighting revealed behaviorally and along the auditory hierarchy.

### A Mechanistic Model Unifying the Start–End Effect Across Neural and Behavioral Levels

To determine whether the observed start–end effects arise from fundamental biophysical constraints on neural coding, we developed a two-stage computational model (Fig. 8). The model incorporates two core principles: dynamic range saturation at the single-neuron level and temporal integration at the population level.

**Figure 8.**
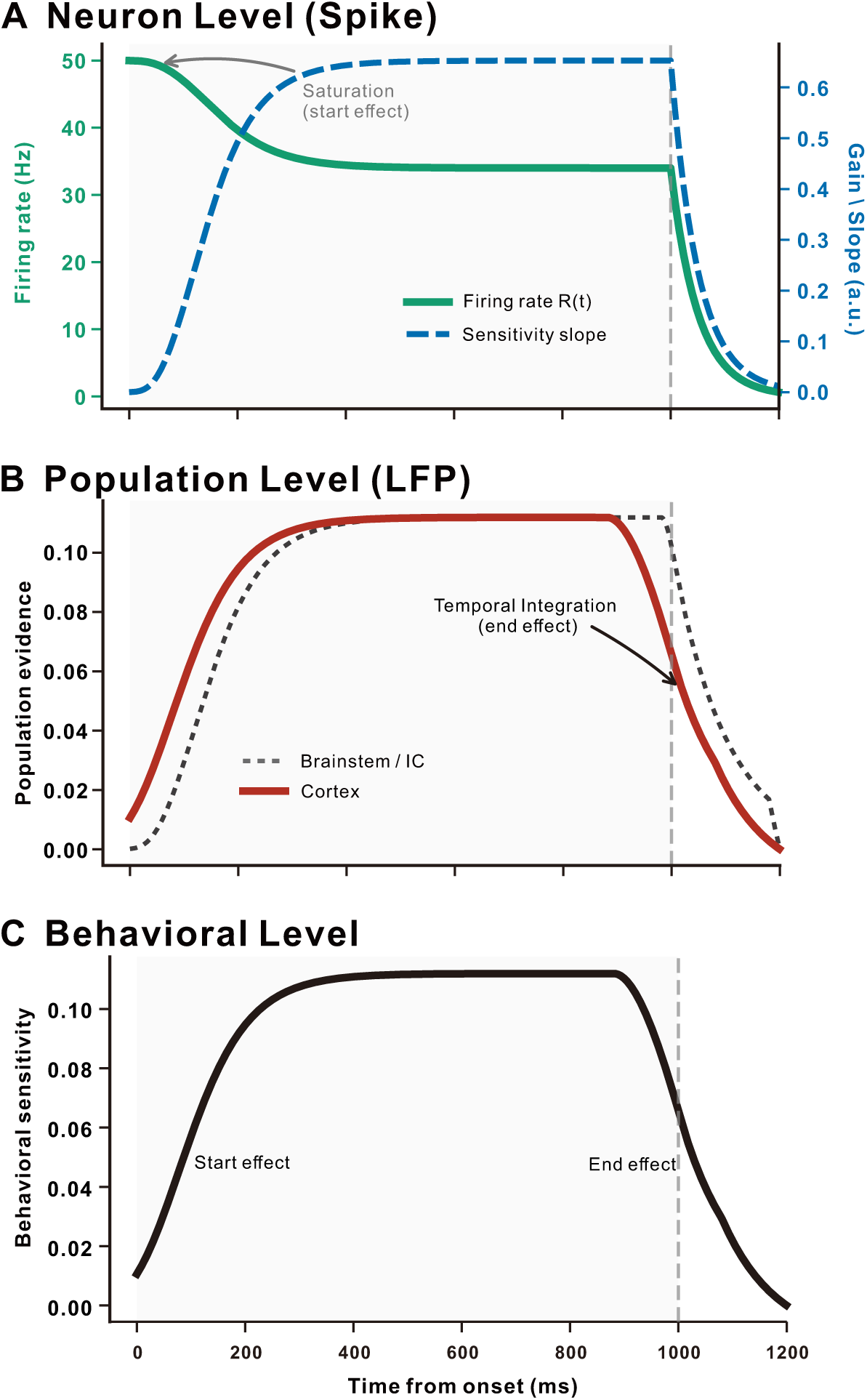
A mechanistic neural model explains the start–end effect through dynamic range saturation and temporal integration. **(A)** Saturation limits onset sensitivity at neuron level (Spike). Simulation of single-neuron dynamics using a Naka-Rushton nonlinearity with adaptation. The green line represents the firing rate R(t), and the blue dashed line represents the instantaneous encoding gain (sensitivity slope). At stimulus onset, the high gain required for detection drives the neuronal response into a saturation plateau. This results in a paradox where the firing rate is maximal, but the encoding gain is compressed to near-zero, providing a biophysical basis for the “start effect” observed at the single-unit level. Both signals exhibit a physiological decay after stimulus offset (1000 ms). **(B)** Integration window creates the end effect at population level (LFP). Comparison of evidence accumulation across hierarchical levels. The black dotted line represents the instantaneous evidence (proxy for Brainstem/IC responses), which tracks the stimulus duration with minimal integration (simulating a short window, e.g., 20 ms). The red solid line represents the integrated cortical evidence, calculated by applying a sliding temporal integration window (∼120 ms) to the instantaneous input. As the integration window crosses the stimulus boundary (vertical dashed line), the inclusion of post-stimulus silence (or decay) naturally dilutes the average evidence, causing the sensitivity to drop prior to stimulus termination (“end effect”). **(C)** Unified sensitivity profile at behavioral level. The final simulated behavioral sensitivity recapitulates the empirical inverted U-shaped profile. The model demonstrates that the temporal weighting of sensory evidence is shaped by two fundamental biophysical constraints: dynamic range saturation at the onset (limiting prospective encoding) and temporal integration windows at the offset (limiting retrospective accumulation).

First, to simulate spike-level responses, we modeled neuronal firing rates using a Naka-Rushton nonlinearity with adaptation dynamics (see Methods for details). At stimulus onset, the system operates with high gain to ensure detection, driving neuronal responses into a saturation regime. Our simulation shows that although firing rate peaks at onset, the instantaneous encoding gain (sensitivity slope) paradoxically drops to near zero (Fig. 8A). This confirms that the behavioral “start effect” results from physiological saturation, a fundamental limitation in neural coding that prioritizes detection over discrimination.

Second, to simulate population-level evidence accumulation (corresponding to cortical LFPs), we applied a temporal integration window (∼120 ms) to the instantaneous output. For the brainstem/IC stage (modeled with a 20 ms integration window), the sensitivity profile tracks stimulus duration with minimal decay (Fig. 8B, dotted line). In contrast, at the cortical stage, the integration window spans the stimulus boundary. As the sliding window crosses the stimulus offset, the inclusion of post-stimulus silence naturally dilutes accumulated evidence, resulting in a premature decline in sensitivity (Fig. 8B, solid line).

Crucially, this simple mechanistic framework—combining onset saturation and offset integration decay—recapitulates the inverted U-shaped sensitivity profile observed in human behavior (Fig. 8C). These results demonstrate that the start–end effect is not a strategic cognitive suppression, but rather an emergent property of a system constrained by bandwidth (at onset) and temporal integration (at offset).

## Discussion

Our results demonstrate that continuous sounds are not processed with uniform sensitivity over time. Instead, human perception and neural activity in humans and rats exhibit a robust start–end effect: sensitivity to brief local changes is attenuated near sound onset and offset and maximal in the middle of the stimulus (Figs. 1–3). Behaviorally, this yields an inverted U-shaped dependence of detection performance on change timing across auditory features and even in vision (Fig. 1; Supplementary Figs. 1–2). Neurally, the same temporal weighting profile emerges across the auditory hierarchy, culminating in a pronounced population-level signature in cortical field potentials, laminar activity, and oscillations (Figs. 2–7). Moreover, a simple, biophysically motivated computational model combining onset saturation with offset-related integration decay (Fig. 8) recapitulates this profile across levels, linking the behavioral and neural start–end effects to basic coding constraints. Together, these findings identify temporal boundary gating as a candidate organizing principle by which sensory processing reweights local evidence both prospectively (following onset) and retrospectively (approaching offset), favoring mid-epoch information.

### Non-uniform temporal weighting as an organizing principle

Across a coordinated set of behavioral and physiological experiments, we consistently observed marked temporal non-uniformity in the encoding of continuous sounds. In humans, local change detection was poorest near stimulus onset and offset and maximal in the mid-portion of the tone (Fig. 1), and this pattern generalized across acoustic features (frequency and amplitude) and into vision (Supplementary Figs. 1–2). In rats, auditory cortical ECoG and single-unit recordings revealed an analogous temporal weighting profile: changes occurring in the middle of the sound evoked robust neural responses, whereas boundary-proximal changes were strongly attenuated (Figs. 3–5). The convergence of psychophysics, EEG, ECoG, and laminar electrophysiology across species suggests that the start–end effect reflects a conserved property of temporal processing in the auditory system, rather than an idiosyncrasy of a particular paradigm (Large and Jones 1999, Naatanen, Paavilainen et al. 2007, Nobre and van Ede 2018).

A key aspect of this principle is its generality. The start–end profile is not limited to frequency perturbations but extends to amplitude perturbations in the auditory domain (Supplementary Fig. 1) and to local changes in a visual motion trajectory (Supplementary Fig. 2), suggesting that similar temporal weighting rules may be implemented across modalities (Oberfeld, Fischenich et al. 2024). Across these settings, the brain appears to apply a simple rule: prioritize the more stable, information-rich middle segment of an event while down-weighting transitional boundaries, where contextual resets and high variability can compromise reliability (Naatanen, Paavilainen et al. 2007, Matthews and Meck 2016). Conceptually, temporal boundary gating complements and extends existing views of temporal attention and predictive coding by showing that the “precision” assigned to local evidence is not constant within an event, but systematically shaped by its internal boundaries (Friston 2005, Feldman and Friston 2010, Clark 2013). Consistent with this principle, our mechanistic model (Fig. 8) shows that such non-uniform weighting can arise from generic coding constraints—onset saturation and finite integration windows—without requiring any task-specific strategy.

### Links to temporal attention, event boundaries, masking and prediction

Our behavioral data provide a psychological bridge between temporal boundary gating and theories of attention, event segmentation, and predictive processing. Detection thresholds were highest for perturbations near onset and offset and lowest in the middle of the tone (Fig. 1E–F), consistent with models in which temporal attention is allocated non-uniformly over time (Large and Jones 1999, Matthews and Meck 2016, de Lange, Heilbron et al. 2018). At sound onset, attention is rapidly oriented to the new event, but the system is still establishing a stable internal model of its temporal and spectral structure. In this regime, prediction errors are low precision and fine-grained local changes are poorly encoded (Friston 2005, Feldman and Friston 2010, Clark 2013). As the sound continues, a stable temporal template or rhythm emerges, increasing the precision of prediction errors and boosting sensitivity to mid-stimulus changes (Large and Jones 1999, Nobre and van Ede 2018). Near offset, attention and internal models begin to shift toward upcoming events or response preparation, decreasing the precision assigned to late evidence and thereby reducing sensitivity again (Matthews and Meck 2016, de Lange, Heilbron et al. 2018).

Onsets and offsets are also canonical event boundaries, known to segment continuous narratives into discrete episodes and to impose transient processing costs (Bregman 1990, Zacks, Speer et al. 2007). In our framework, these boundaries act as strong “reset” signals that consume representational and mnemonic resources, effectively reducing the gain on local evidence sampled near those transitions. At more peripheral levels, the same pattern can be understood in terms of forward and backward masking: energy at the start and end of the sound masks nearby perturbations, whereas the mid-portion provides a stable reference and lower masking (Oxenham 2001, Moore 2013). Thus, temporal boundary gating can be viewed as the emergent outcome of interacting processes—time-varying attention, masking, event segmentation, and predictive coding—that jointly determine how sensory evidence is weighted across an event (Naatanen, Paavilainen et al. 2007, Liu, Tsunada et al. 2015, de Lange, Heilbron et al. 2018).

### Circuit and population mechanisms of the start–end effect

Our pathway-wide recordings reveal how temporal boundary gating is implemented at the circuit level. Population analyses across IC, MGB, and AC showed a robust start effect: early-sequence responses to local changes were consistently suppressed relative to mid-sequence responses (Fig. 5). The magnitude of this onset-locked suppression increased along the hierarchy—weakest in IC, intermediate in MGB, and strongest in AC—consistent with cumulative adaptation and sharpening by thalamocortical and intracortical circuitry (Wehr and Zador 2003, Ulanovsky, Las et al. 2004, Wehr and Zador 2005, Malmierca, Cristaudo et al. 2009, Antunes and Malmierca 2011, Pérez-González, Hernández et al. 2012, Yaron, Hershenhoren et al. 2012, Kopp-Scheinpflug, Sinclair et al. 2018). This progression parallels known increases in intrinsic and synaptic timescales from midbrain to cortex (Hasson, Yang et al. 2008, Murray, Bernacchia et al. 2014, Chaudhuri, Knoblauch et al. 2015), which lengthen the window over which recent stimulation shapes subsequent responses.

Mechanistically, the start effect likely reflects onset-conditioned encoding: strong onset-evoked excitation is followed by rapid suppression that transiently reduces sensitivity to nearby changes. Forward suppression and short-term synaptic depression at thalamocortical and intracortical synapses, together with fast feedforward inhibition, are well-established contributors to this phenomenon (Wehr and Zador 2003, Wehr and Zador 2005, Natan, Briguglio et al. 2015). Superimposed on this, stimulus-specific adaptation (SSA) further suppresses responses to predictable or repeated patterns and recovers over hundreds of milliseconds (Ulanovsky, Las et al. 2004, Malmierca, Cristaudo et al. 2009, Antunes and Malmierca 2011, Taaseh, Yaron et al. 2011, Pérez-González, Hernández et al. 2012, Yaron, Hershenhoren et al. 2012). A simple picture emerges: a strong onset response is followed by suppression and adaptation that gradually recover, yielding the monotonic rise and plateau of local-change responses from early to mid positions in IC→MGB→AC (Fig. 5D–E).

In contrast, the end effect is less prominent and more heterogeneous at the single-unit level. While some AC neurons exhibit reduced firing near the end of the sequence, end-related changes in spike rate are weaker and more variable than start-related suppression overall (Fig. 5D–E). However, our laminar LFP recordings reveal that the full inverted U-shaped profile is clearly expressed at the population level in cortex. In A1, ΔRM in the thalamorecipient granular layer shows minimal responses for early and late changes and a pronounced maximum for mid-sequence changes, with parallel but weaker modulation in supra- and infragranular layers (Fig. 6D–E). Time–frequency analyses further show that alpha, beta, and gamma power are maximally enhanced by mid-stimulus changes and attenuated near both onset and offset across layers (Fig. 7B–C). Because LFPs and band-limited power primarily reflect synchronized population activity rather than mean firing alone (Ray and Maunsell 2011, Buzsáki and Wang 2012, Einevoll, Kayser et al. 2013), these findings indicate that the end effect is implemented largely through changes in population synchrony and laminar-specific network dynamics, rather than a simple decrease in average firing.

End-related responses themselves likely involve slower and more diverse mechanisms than onset suppression, including disinhibition, post-inhibitory rebound (e.g., via T-type Ca²⁺ channels), and specialized offset-tuned circuits in subcortical and cortical stations (Scholl, Gao et al. 2010, Aubie, Sayegh et al. 2012, Kopp-Scheinpflug, Sinclair et al. 2018, Solyga and Barkat 2021, Song, Xu et al. 2024). These processes unfold over tens to hundreds of milliseconds and are distributed across layers and cell types, which explains why they are robust in LFP and oscillatory measures (Figs. 6–7), yet muted and heterogeneous in short-window spike-rate metrics. In this sense, the cortical end effect can be viewed as a retrospective temporal weighting process: as the system approaches stimulus termination, cortical networks integrate recent input over relatively long windows and adjust the gain on late evidence in anticipation of the upcoming boundary (Buzsáki and Wang 2012, Murray, Bernacchia et al. 2014, Chaudhuri, Knoblauch et al. 2015). Consistent with this interpretation, the population-level start–end asymmetry is reproduced by our model (Fig. 8).

### Temporal integration windows and bidirectional gating

Our findings suggest that the brain’s non-uniform sampling of continuous events—prioritizing the middle while suppressing the boundaries—is a normative solution to the trade-off between detection and discrimination.

At the computational level, local change detection is supported by temporal integration windows that extend both backward and forward in time around the change. Effective detection requires integrating evidence across this window, retrospectively and prospectively. When a change occurs near onset, the strong onset response intrudes into the prospective portion of the window; when a change occurs near offset, impending termination and offset-related dynamics intrude into the retrospective portion. In both cases, boundary events truncate the effective integration window for the local perturbation, thereby reducing the amplitude of behavioral and neural change responses and producing the observed start–end effect across psychophysics and physiology (Figs. 1–7).

Our mechanistic model provides a concrete explanation for how such bidirectional gating can arise from generic coding constraints (Fig. 8). We show that the “start effect” arises from a dynamic range trade-off: to rapidly detect the onset of an event, the system must maximize gain, which inevitably drives neurons into saturation (Albrecht and Hamilton 1982, Carandini and Heeger 2011). This saturation effectively “blinds” the system to prospective information, preventing the high-fidelity encoding of local details immediately following detection. Conversely, the “end effect” reflects the constraints of retrospective integration. While subcortical circuits (IC) track instantaneous inputs with high temporal fidelity, cortical circuits accumulate evidence over longer timescales (∼120 ms) to stabilize perception (Hasson, Yang et al. 2008, Murray, Bernacchia et al. 2014, Sabat, Gouyette et al. 2025). Our model demonstrates that stimulus termination creates a “moving average decay,” where the cessation of afferent input dilutes the evidence available within the cortical integration window. This supports the hypothesis that event boundaries disrupt the retrospective accumulation of sensory evidence (Zacks, Speer et al. 2007). Together, these mechanisms establish a hierarchical “boundary gating” architecture: physiological saturation limits sensitivity at the start (hardware limit), while integration dynamics limit sensitivity at the end (computational limit).

This bidirectional gating challenges models that treat continuous auditory processing as temporally uniform. Instead, our data support a view in which the brain implements a strong, spike-level “start gate” that transiently down-weights early evidence, combined with a more diffuse, oscillation-based “end gate” that retrospectively evaluates late evidence in light of impending stimulus termination. Temporal boundary gating thus provides a mechanistic link between adaptation, predictive coding, and event segmentation, embedding them within a common temporal-integration framework.

### Implications and outlook

Beyond its conceptual contribution, the start–end effect has several practical implications. In engineered systems, perceptually inspired audio and video compression schemes (e.g., MP3, MPEG) could exploit temporal non-uniformity by allocating fewer bits to onset and offset segments while preserving fidelity in the mid-epoch (Painter and Spanias 2000). Computational models of attention and prediction could incorporate explicit time-varying precision profiles to better capture neural and behavioral data in continuous tasks (Friston 2005, Feldman and Friston 2010, Clark 2013, de Lange, Heilbron et al. 2018). In neurotechnology, brain–computer interfaces and decoding algorithms might improve robustness by down-weighting boundary-locked variability and emphasizing mid-epoch signals (Wolpaw, Birbaumer et al. 2002, Banno, Lestang et al. 2020). For hearing aids and cochlear implants, emphasizing mid-utterance cues—where our data indicate maximal sensitivity—may yield perceptual gains in noisy, real-world environments, in which onsets and offsets are often masked or distorted (Oxenham 2001, Moore 2013).

In summary, our work identifies and mechanistically dissects a robust, conserved start–end effect in sensory processing, and introduces temporal boundary gating as a core candidate principle of how the brain weights information within continuous events. By linking behavior, macroscopic signals, and laminar circuit dynamics across species, these findings provide a framework for understanding how the auditory system—and potentially other sensory systems—allocates computational resources across time to support stable perception in a dynamic world.

## Methods

### Participants and Surgery

#### EEG Experiment Participants

The study involved psychological experimental sessions and EEG recording experimental session, all conducted in accordance with the Declaration of Helsinki. For psychological experiment, *Frequency Session 1* included 19 volunteers (13 males and 6 females; aged 21–34 years) with normal hearing. Out of these, 14 volunteers participated in *Frequency Session 2* and 3, and 11 volunteers participated in *Amplitude Session 4* and *5*. There are also 10 volunteers take part in *Visual Session 6*. Besides, a total of 30 volunteers (14 males and 16 females; aged 20–35 years) took part in EEG record experiment. The study protocol was approved by the Institutional Review Board (IRB-20230131-R), and informed consent was obtained from all participants prior to their inclusion.

#### ECoG surgery

Five adult male Wistar rats (weighing 280 - 340 g and aged 9 - 12 weeks), with healthy external ears, were selected for the implantation of the headpost and ECoG array. Anesthesia was administered using pentobarbital sodium (40 mg/kg), with atropine sulfate (0.05 mg/kg, s.c.) given 15 minutes prior to the procedure to reduce tracheal secretion. Xylocaine (2%) was applied generously to the incision site to minimize pain. A head fixation bar was attached to the top of the skull using dental cement and six titanium screws. A craniotomy exposed a 4.5 mm × 5 mm area over the auditory cortex on the left side, with the dura mater remaining intact. The ECoG array (KD-PIFE, KedouBC, China), comprising 32 electrodes in a 6×6 grid with 1 mm spacing, was implanted. The array covered approximately an area of 5 mm × 5 mm, encompassing the entire auditory cortex (Fig. 3A). Post-implantation, the ECoG array was covered with artificial brain gel, and the bone flap was repositioned over the surgical site. The area was sealed with dental cement to ensure closure and protection. The postoperative recovery period lasted seven days, during which anti-edema medications and antibiotics were administered, and the animals’ weights were monitored daily to ensure their well-being. This part of the study was approved by the Animal Subjects Ethics Committees of Zhejiang University (ZJU20210078).

#### Extracellular recording surgery

Three adult male Wistar rats (weighing 280 - 340 g and aged 9 - 12 weeks) were used for extracellular electrode recordings, undergoing a procedure similar to the ECoG surgery for head post stabilization. The surgical and recording procedures were performed as previously described (Zhai, Sun et al. 2019, Zhai, Auksztulewicz et al. 2020, Song, Zhai et al. 2021, Gong, Zhai et al. 2022). Following surgery, the left lateral and dorsal surfaces of the skull were exposed. Once cleaned and dried, reference landmarks were established at the bregma and on the left lateral skull for precise electrode positioning. The target areas for electrode recordings included the left primary auditory cortex (A1), the left medial geniculate body (MGB), and the left inferior colliculus (IC). To prevent tissue overgrowth, a thin layer of dental cement was applied over the skull above these areas, and a protective wall was constructed. A handheld micro-drill was used to create a small opening (approximately 2 mm × 2 mm) at the designated target site, which was subsequently sealed with brain gel. On the recording day, the dura mater was carefully punctured, allowing the electrode to be vertically inserted into the target area, guided by the pre-established landmarks.

### Stimuli and Experimental Procedures

Detailed stimulus parameters for human psychophysical experiments are summarized in *Table 1*. Electrophysiological experiment stimuli parameters, including EEG, ECoG, and extracellular recordings, are presented separately in *Table 2*.

#### Psychological Experimental Procedures

##### Frequency Session 1: frequency-gradient manipulation (*Fig. 1D*)

In this session, participants performed a delayed match-to-sample task involving 500 ms pure tones at 1 kHz. Each trial started with the presentation of a 500 ms reference tone, followed by a 500 ms silent interval, then a second 500 ms test tone. The test tone contained a ∼20 ms local frequency perturbation (approximately 20 cycles of the 1 kHz carrier) embedded at one of three temporal positions: 25 ms, 250 ms, or 475 ms (5%, 50%, 95% of duration). The magnitude of the frequency perturbation varied across five levels (1%, 2%, 3%, 4%, and 6%), plus a no-change control condition (1 kHz). Each condition was repeated 40 times, resulting in 16 different auditory stimuli randomly selected for presentation. Participants assessed whether the two sounds were identical, using the left arrow key for differing sounds and the right arrow key for identical sounds.

##### Frequency Session 2: temporal-position manipulation (500 ms) (*Fig. 1A*)

In this session, participants performed a change detection task using 500 ms pure tones at 1 kHz. Within each tone, a ∼20 ms local frequency perturbation (20 cycles) was introduced at one of eleven temporal positions, corresponding to 25, 50, 75, 100, 150, 250, 350, 400, 425, 450, or 475 ms (5%, 10%, 15%, 20%, 30%, 50%, 70%, 80%, 85%, 90%, and 95% of the tone duration). The perturbation magnitude was fixed at a level previously established to yield approximately 60% detection accuracy at 250 ms in *Frequency Session 1*. Each temporal condition was repeated 40 times, resulting in 12 unique auditory stimuli (including no-change control), presented in pseudorandom order. Participants determined whether a change occurred during each sound, pressing the left keyboard key for a change and the right key if no change was perceived.

##### Frequency Session 3: temporal-position manipulation (1000 ms) (*Fig. 1A*)

This session replicated the design of *Frequency Session 2*, except that the pure tones lasted 1000 ms. The same ∼20 ms local frequency perturbation was introduced at eleven temporal positions corresponding to 50, 100, 150, 250, 350, 500, 650, 750, 850, 900, or 950 ms (5%, 10%, 15%, 25%, 35%, 50%, 65%, 75%, 85%, 90%, and 95% of the tone duration). Each condition was repeated 40 times, yielding 12 unique stimuli (including no-change control) presented in random sequence. Participants performed the same change detection task as in *Frequency Session 2*.

##### Amplitude Session 4: amplitude-gradient manipulation (Supplementary Fig. 1A)

This session employed a delayed match-to-sample task analogous to *Frequency Session 1*. Participants first heard a 500 ms pure tone at 1 kHz, followed by a 500 ms silent interval, and then a second 500 ms tone. The second tone contained a ∼20 ms local amplitude perturbation (20 cycles at 1 kHz), implemented as an increase in amplitude of one of five magnitudes: 2%, 3%, 4%, 5%, or 6%. This perturbation was embedded at one of three temporal positions corresponding to 25 ms, 250 ms, or 475 ms (5%, 50%, and 95% of the total tone duration). Each condition was repeated 40 times, resulting in 16 unique stimuli (including no-change control) presented in randomized order. The carrier frequency remained constant at 1 kHz throughout, ensuring that only amplitude varied locally. Participants indicated whether the second tone was identical to the first via designated response keys.

##### Amplitude Session 5: amplitude-position manipulation (Supplementary Fig. 1C)

Using a change detection task similar to *Frequency Session 2*, this session presented 500 ms pure tones at 1 kHz containing a ∼20 ms local amplitude perturbation at one of eleven temporal positions: 25, 50, 75, 100, 150, 250, 350, 400, 425, 450, or 475 ms (5%, 10%, 15%, 20%, 30%, 50%, 70%, 80%, 85%, 90%, and 95% of tone duration). The amplitude perturbation magnitude was fixed to match the level yielding approximately 60% detection accuracy at 250 ms in *Amplitude Session 4*. Each temporal condition was repeated 40 times, resulting in 12 unique stimuli (including no-change) presented in pseudorandom order. Participants judged whether an amplitude change occurred within the tone by pressing designated response keys. The carrier frequency remained constant at 1 kHz.

##### Visual Session 6 (Supplementary Fig. 2A)

This visual change detection task involved a black vertical bar (13 pixels wide) moving horizontally across the central part of the screen with a resolution of 2560 × 1440 pixels. The bar moved from 30% to 70% of the screen width (covering a total distance of 512 pixels) at a speed of approximately 30 pixels per frame, completing its movement in 17 frames. During the movement, the bar changed to gray for two frames at relative positions of 50, 100, 150, 250, 500, 750, 850, 900, or 950 ms (5%, 10%, 15%, 25%, 50%, 75%, 85%, 90%, and 95% along its path). Each of the 10 conditions—including 9 color-change positions and a control condition without color change—was repeated 40 times. Participants were instructed to judge whether a color change occurred by pressing the left arrow key for changes and the right arrow key for no changes.

#### Human EEG Experimental Procedures

##### Frequency Session: temporal-position manipulation (Fig. 2)

Similar to the *psychological experiment*, *Frequency Session 3*, 1000 ms pure tones at 1 kHz contained local frequency perturbations lasting ∼20 cycles. The magnitude of frequency increase was fixed at 8%, determined through a stepwise procedure: initial perturbations around the threshold level used in psychological experiments did not elicit any detectable neural change responses. Consequently, the perturbation magnitude was gradually increased—testing 4%, 6%, and finally 8%—until reliable change-related neural activity was consistently observed. These perturbations were embedded at eleven temporal positions: 50, 100, 150, 200, 250, 500, 750, 800, 850, 900, or 950 ms (5%, 10%, 15%, 20%, 25%, 50%, 75%, 80%, 85%, 90%, and 95% of the tone duration). Each condition (no-change condition included) was presented 45 times in random order. Although participants were not required to perform any behavioral task during this session, continuous electroencephalogram (EEG) recordings were acquired to assess neural responses to the frequency changes.

Auditory stimuli were generated by MATLAB, and all frequency perturbations were implemented in a phase-continuous manner. Specifically, frequency changes were introduced only at zero-crossing points of the carrier waveform, such that transitions from 1 kHz to the perturbed frequency (e.g., 1.02 kHz) occurred at the end of an integer number of carrier cycles, and the return to 1 kHz likewise occurred after completion of an integer number of cycles of the perturbed frequency. This ensured that frequency perturbations did not introduce phase discontinuities or temporal jitter. The same phase-continuous procedure was used for all local frequency manipulations throughout the study.

And these experiments above were conducted in a sound-proof room, where participants maintained a static head position while listening to auditory stimuli and responded via keyboard presses. Auditory stimuli were delivered using a Golden Field M23 sound player, powered by a Creative AE-7 Sound Blaster, with a sampling rate of 384 kHz. Sound delivery was controlled using Psychtoolbox 3 in MATLAB.

#### Rat ECoG Experimental Procedures

Three passive listening sessions, identical to those used in human EEG studies, were adapted for use with rats. ECoG and Extracellular recording experiments took place in a sound-proof room, with stimuli played contralaterally to the recording site in rats. These acoustic stimuli were digitally generated using a computer-controlled Auditory Workstation (RZ6, Tucker-Davis Technologies) with a sampling rate of 97656 Hz and delivered through magnetic speakers (MF1, TDT) within the TDT systems.

##### Frequency Session 1: temporal-position manipulation (Fig. 3 and Supplementary Fig. 3)

This session involved auditory stimuli comprising 14 conditions, each consisting of complex sound mixtures at frequencies of 4 kHz, 8 kHz, 16 kHz, and 32 kHz. Each sound lasted 1000 ms, with local frequency changes occurring at specific temporal positions: no-change, 25, 50, 75, 100, 150, 250, 500, 750, 850, 900, 925, 950, or 975 ms (2.5%, 5%, 7.5%, 10%, 15%, 25%, 50%, 75%, 85%, 90%, 92.5%, 95%, and 97.5% of the total duration). For each frequency component, a 3.6% increase was applied lasting 4 ms (corresponding to 16 cycles for 4 kHz or 32 cycles for 8 kHz, etc.). The choice of this 3.6% increment was empirically determined during ECoG recordings by testing frequency changes at the 50% position within the sound. A series of frequency changes at the 50% position within the sound were tested, incrementally increasing from 2% to 8% in steps of 0.4%. Based on these tests, 3.6% was selected as the frequency change magnitude that corresponded approximately to a 60% saturation level in the neural response. Although slight individual variability existed among rats, this 3.6% frequency change value was consistently similar across rats and thus was adopted as the standard parameter. In the control condition, no frequency change was applied. Each stimulus condition was presented 40 times in a random order.

##### Amplitude Session 2: temporal-position manipulation (Supplementary Fig. 4)

This session included auditory stimuli consisting of 12 conditions, each composed of complex sound mixtures at 4 kHz, 8 kHz, 16 kHz, and 32 kHz. Each stimulus lasted 500 ms, with intensity changes occurring at specific time points corresponding to 25, 50, 75, 100, 150, 250, 350, 400, 425, 450 or 475 ms (5%, 10%, 15%, 20%, 30%, 50%, 70%, 80%, 85%, 90%, and 95% of the total duration). For each frequency component, the amplitude was increased by 36% over a 4 ms window (resulting in 16 cycles for 4 kHz and 32 cycles for 8 kHz, etc.). The amplitude increment of 36% was empirically determined during the ECoG recordings by testing intensity changes at the 50% position within the stimulus. A range of increments from 20% to 80% was applied, and the variability of the neural responses was computed over a 150 ms window following the change onset. The 36% increment was chosen as it corresponded approximately to a 60% saturation level in the evoked responses. Despite slight individual differences among rats, this 36% amplitude change was consistent across subjects and therefore used as the standard parameter. In the control condition, no amplitude changes were applied. Each stimulus condition was repeated 40 times and presented randomly.

#### Rat Extracellular recording experimental procedures

Firstly, to characterize basic neuronal response properties, we assessed the frequency response area (FRA) and determined characteristic frequency (CF) by presenting randomized pure tones across a range of frequencies and intensities at the contralateral position relative to the recording site. Subsequently, a local change detection paradigm was employed to investigate neural sensitivity to stimulus alterations.

##### Frequency Session: frequency-position manipulation (Fig. 4-5 and Supplementary Fig. 5-6)

The auditory stimulus in this session consisted of 14 conditions, with each sound being a complex harmonic tone comprised of frequencies at 4 kHz, 8 kHz, 16 kHz, and 32 kHz, with a total duration of 1000 ms. Similarly to *rat ECoG experiment, Frequency Session 1*, frequency changes occurred at specific positions: 25, 50, 75, 100, 150, 250, 500, 750, 850, 900, 925, 950, or 975 ms (2.5%, 5%, 7.5%, 10%, 15%, 25%, 50%, 75%, 85%, 90%, 92.5%, 95%, and 97.5% of the duration). For each frequency, a 3.6% increase was applied for 4 ms, corresponding to 16 cycles for 4 kHz and 32 cycles for 8 kHz, among others. The 3.6% frequency increase was based on the threshold frequency difference known to elicit a change response same as *ECoG Session 1*. In the control condition, the frequency remained unchanged. Each sound condition was repeated 40 times and presented in a random order.

Sound intensity was uniformly calibrated to 60 dB SPL to ensure consistency across experiments involving both human participants and animal subjects. Calibration was performed using a ¼-inch condenser microphone (Brüel & Kjær 4954, Nærum, Denmark) and a PHOTON/RT analyzer (Brüel & Kjær, Nærum, Denmark).

### Data acquisition and pre-processing

#### EEG recording

Electroencephalogram (EEG) signals were recorded using a 64-channel electrode cap based on the international 10-20 system (NeuSen W series, Neuracle, China) combined with a wireless amplifier (NeuSen W series, Neuracle, China). Five additional electrodes monitored electrooculography (EOG) and electrocardiogram (ECG) signals (‘ECG’, ‘HEOR’, ‘HEOL’, ‘VEOU’, and ‘VEOL’), leaving 59 channels dedicated to EEG analysis. Reference and ground electrodes were placed posterior and anterior to the Fz electrode, respectively. EEG data and trial onset markers were simultaneously acquired at a sampling rate of 1 kHz. Participants sat comfortably in an acoustically shielded room and were instructed to minimize blinking and facial movements to reduce artifacts.

EEG preprocessing followed standard procedures commonly used in electrophysiological analysis and EEG-based classification studies(Tu, Zhang et al. 2025, Xu, Huang et al. 2025). Raw signals were bandpass filtered between 0.5 and 40 Hz and segmented into epochs spanning -1 to 2 seconds relative to trial onset. Independent component analysis (ICA) was applied to remove ocular artifacts. Baseline correction was performed by subtracting the mean signal within the -200 to 0 ms pre-stimulus interval from each epoch. Artifact rejection employed a relative thresholding approach: samples exceeding mean ± 3 × standard deviations (SD) were labeled as “bad samples.” Trials with more than 20% bad samples were classified as “bad trials”, and channels with over 10% bad trials were identified as “bad channels”. Bad channels were excluded after comprehensive assessment across all channels, and bad trials were removed for all channels based on the remaining good channels’ data.

Event-related potentials (ERPs) were obtained by averaging epochs for each condition, channel, and subject. To mitigate inter-channel variability, ERP amplitudes were normalized by each channel’s standard deviation within subjects before group-level analyses.

#### ECoG recording

The ECoG recording procedure involved placing a reference wire from the ECoG array into the subdural space, with the ground wire firmly connected to a titanium screw located at the skull’s front. Lead wires from the ECoG array were attached to micro connectors (ZIF-Clip headstage adapters; Tucker-Davis Technologies, TDT, Alachua, FL). The recorded signals were amplified using an RZ5 amplifier (TDT), digitally sampled at a 12 kHz rate, and stored on hard drives for subsequent detailed analysis.

Raw electrocorticography (ECoG) data were down-sampled to 600 Hz and subjected to a 50 Hz notch filter to eliminate power line interference. The continuous signals were segmented into epochs of 2 seconds duration, except for signals lasting 500 ms, which were segmented into 1.5-second epochs. Epochs spanned from -0.5 seconds before trial onset to 0.5 seconds after trial offset. Baseline correction was applied to each epoch to account for pre-stimulus signal variations. Channels or trials identified as problematic during acquisition were excluded from subsequent analyses. The preprocessing procedures mirrored those applied to EEG data to maintain methodological consistency.

#### Extracellular and LFP recording

For extracellular and local field potentials (LFP) recordings, Version 3A Neuropixels silicon probes (Imec, Leuven, Belgium; available at https://github.com/cortex-lab/neuropixels/wiki/About_Neuropixels) were employed during each session. Track positions were meticulously aligned using pre-established markers to ensure precision. A flexible ground wire was soldered to the probe’s printed circuit board (PCB), with the ground and reference contacts shorted. Throughout the experimental procedures, this ground wire was connected to a titanium screw at the frontal skull area. The probes were carefully inserted vertically through 0.3−0.5 mm slots in the dura, meticulously avoiding visible blood vessels to minimize tissue damage. The insertion process was facilitated by a single-axis motorized micromanipulator, controlled remotely from outside the soundproof chamber. For A1, the average insertion depth was approximately 2.0 ± 0.1 mm (n = 12); for MGB, it was about 7 ± 0.5 mm (n = 14); and for IC, approximately 6.1 ± 0.4 mm (n = 12) below the brain surface, consistent with the standard rat brain atlas (Paxinos and Watson 2014). White noise stimuli were utilized to confirm the presence of auditory responses.

Neuronal signals were acquired using the Neuropixels PXIe control system (Imec, Leuven, Belgium) at sampling rates of 30 kHz for spike data and 2.5 kHz for LFP. Spike sorting was performed offline with the Kilosort algorithm and manually validated using Phy (Pachitariu, Steinmetz et al. 2016). For each cluster, we calculated the mean firing rate during the pre-stimulus period. Three criteria were applied to identify auditory-responsive neurons: (1) the number of neurons showing a significant increase in firing rate during the onset window (0–200 ms) compared to the baseline window (−200 to 0 ms before onset) was assessed using a paired t-test; (2) post-stimulus firing rates exceeding three times the standard deviation of the baseline firing rate; and (3) neuronal latency was computed using a custom MATLAB function employing a Poisson statistical model, with responsive neurons required to have a non-empty latency result. Clusters meeting all these criteria were classified as auditory-responsive and included in subsequent analyses. A total of 671 neurons from A1, 632 neurons from MGB, and 940 neurons from IC were selected accordingly.

Recordings targeting the auditory cortex focused exclusively on the A1 region of the left hemisphere, identified based on the tonotopic gradient of characteristic frequencies. After the recording sessions, the Neuropixel probes were coated with DiI prior to insertion into the medial geniculate body (MGB) and inferior colliculus (IC) to verify accurate probe placement (Harpaz, Jankowski et al. 2021). Following recordings, rats were transcardially perfused under a fume hood.

### Data Analysis and Statistics

The offline processing of EEG, ECoG, LFP and extracellular data were conducted using MATLAB R2023b (MathWorks) and the FieldTrip toolbox (Oostenveld, Fries et al. 2011).

#### Statistical tests

Various statistical analyses were performed to evaluate the data. These included t-tests, ANOVAs, and non-parametric tests, with specific methods indicated parenthetically where applicable.

For example, to determine whether detection effects were linked to stimulus onset or offset (Fig. 1C), detection ratios were calculated at each change position for both stimulus durations (500 ms and 1000 ms) in each participant (n = 14). Wilcoxon signed-rank tests were used to compare detection ratios at corresponding positions between the two durations. This non-parametric test assessed whether detection performance significantly differed when aligned to stimulus onset or offset.

#### Psychological threshold assessment

To evaluate psychological thresholds, the change detection ratio for each group was calculated by dividing the number of trials in which subjects pressed the left arrow key (indicating change detection) by the total number of trials within that group. Subjects were excluded from further behavioral analyses if their change detection ratio was greater than 0.3 in the control group or less than 0.6 in the maximum change group. Ultimately, 19 subjects from *psychological Experiment, Frequency Session 1* (Fig. 1E), and 11 subjects from *psychological* Experiment, *Session 4* (Supplementary Fig. 1B) were included. Psychometric functions were fitted to the data using a cumulative Gaussian function (Yu, Dickman et al. 2015, Xu, Zhai et al. 2017):

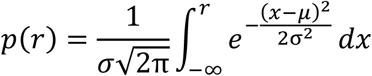

where 𝑝(𝑟) represents the ratio of change detection as a function of local change value. Parameters 𝜇 and 𝜎 represent the mean and standard deviation of Gaussian fit, respectively. The psychological threshold for change detection was defined at 𝑝(𝑟) = 0.6 on the fitted curve. This curve-fitting procedure was performed using the ‘psignifit’ MATLAB toolbox (available at http://bootstrap-software.org/psignifit/).

#### EEG data analysis

##### Permutation test

To compare event-related potentials (ERP) or global field power (GFP) between two conditions at the time-sample level, we employed a two-tailed cluster-based permutation test using the ‘ft_timelockstatistics’ function from the FieldTrip toolbox in MATLAB. The procedure consisted of the following steps: *1)* At each time position and channel, a two-tailed independent-samples t-test was performed to compare Dataset A and Dataset B, yielding a matrix of t-values. *2)* A cluster-defining threshold of p < 0.05 was applied to identify significant time points. Time points exceeding this threshold were designated as candidate significant points. Clusters were then formed by grouping consecutive significant time points and neighboring significant electrodes (for GFP comparisons, clustering was performed only across time points) utilizing a Statistical Parametric Mapping (SPM) labeling algorithm (Thurfjell, Bengtsson et al. 1992). *3)* Condition labels for Dataset A and B were randomly permuted. For each permutation, a new t-value matrix was generated, and clusters were re-formed using the same threshold. The largest cluster-level statistic (e.g., sum of t-values) from each permutation was retained, creating a null distribution. *4)* The observed clusters from step *2)* were compared to the null distribution. A cluster was considered significant if its cluster-level statistic exceeded 95% of the permuted clusters (p < 0.05).

In this study, the permutation test was performed at the inter-subject level on EEG data to define the time window of change response. Based on this identified window, the relative magnitude change (RM) value was calculated to quantify the difference among conditions. Specifically, for the EEG data, Dataset A consisted of GFP data collected under the control condition (*Human EEG Experiment, Frequency Session*) from 30 subjects, recorded across 64 channels. Dataset B comprised GFP data from the same 30 subjects corresponding to the local change response elicited at the central (50%) position out of 11 stimulus locations. A significant time cluster showing difference between the local change and control conditions was identified, which corresponded approximately to 60–220 ms after the local change onset. This time window was then selected for calculating the RM values.

##### Quantification of change response

To quantify the change responses across different positions, we first averaged the ERP data from each channel across trials for each subject. Figure 2A shows the grand-average waveform from an example channel across all subjects. Next, we computed the RM value for each channel to quantify the ERP magnitude within the time window identified by the permutation test. The RM was defined as the root mean square (RMS) of the ERP signal in this window. We then calculated the difference in RM values (ΔRM) between the change and control conditions across positions. Specifically, RM for channel i over the time window [t_1_, t_2_] was calculated as:

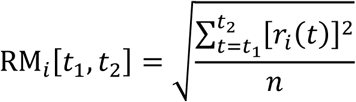

For our EEG data, the selected time window was 60–220 ms following the former description. RM values were calculated for each ERP corresponding to different change positions within this window, as well as for the control condition in the same time window. The ΔRM between change and control conditions was then computed.

To evaluate statistical significance across subjects, an independent-samples t-test was also conducted on the ΔRM values for each positional condition. Channels showing significant differences (p < 0.05) were highlighted in black in Figure 2B. Additionally, we averaged ΔRM values across all channels for each subject to investigate position-dependent changes in the overall change response (Figure 2C). An independent-samples t-test was also performed to assess whether the mean ΔRM across subjects was significantly greater than zero, confirming the presence of a genuine difference across positions.

Specifically, the ΔRM values were normalized by each subject’s maximum ΔRM observed across all recorded positions. The ΔRM at the early time point (e.g., 50 ms, referred to as the start) and at the last position (e.g., 950 ms, referred to as the end) were expressed as ratios relative to this maximum. This normalization controls for individual variability in absolute signal amplitudes and allows for direct comparison between human EEG data and extracellular spike data recorded from rodents, facilitating cross-species analysis of temporal dynamics in neural responses.

#### ECoG data analysis

The permutation test described earlier in this paper was also conducted at the inter-channel level for ECoG to define the change response window. Dataset A represented the GFP data from control condition trials (e.g., *Rat ECoG Experiment, Frequency Session 1*), with each trial providing data across 32 channels. Dataset B consisted of GFP data from the frequency change location at 50%. Based on the significant differences identified by the permutation test between the control and 50% change location groups across various channels, we selected a time window of 0–160 ms following the change onset to calculate RM values. This approach aligns with methods commonly applied in EEG studies.

Using the same root mean square (RM) computation method applied to EEG data, RM values were calculated for each change location within the 0–160 ms window following change onset. These RM values were then compared to those obtained under control conditions in the corresponding time windows. The resulting ΔRM values, representing the difference between change and control conditions, are visualized in the topographical maps shown in Figure 3C. Electrodes exhibiting statistically significant differences (two-tailed independent-samples t-tests) are indicated with black dots on the maps.

Since the ECoG electrode coverage extended beyond the auditory cortex (Figure 3A), we first evaluated each channel’s responses to stimulus onset under the control condition. For each channel, permutation tests were performed comparing neural activity in the pre-stimulus window (−300 to 0 ms) and the post-stimulus window (0 to 300 ms). From the resulting statistical values, we identified the maximum t-value within the early post-stimulus period (0–50 ms) for each channel. Channels were then ranked according to these maximum t-values, and the top 16 channels exhibiting the strongest onset responses were selected for subsequent analyses.

For each trial, ΔRM values from the 16 selected channels were averaged to obtain an RM difference (ΔRM) metric for each change location (Figure 3D).

Additionally, pairwise t-tests with Benjamini-Hochberg false discovery rate (FDR) correction were conducted to determine whether ΔRM differed significantly between change locations, as shown in Figure 3E.

#### Extralcellular data analysis

The data processing of spike recordings followed procedures analogous to those applied for EEG and ECoG data. Specifically, for each neuron and each stimulus condition, firing rates were calculated within a fixed 0–100 ms time window following the onset of the local frequency change at a given stimulus position (Fig. 4B). To quantify neuronal modulation, Δfiring rates (ΔFR) for the local change stimuli were baseline-corrected by subtracting the corresponding firing rate observed in the control stimulus within the same time window. For example, for a stimulus with a frequency change at 25 ms, the firing rate was computed over 25–125 ms and then subtracted by the control condition firing rate in the same 25–125 ms range.

Consequently, each neuron yielded 13 ΔFR values representing modulation relative to control across all local change positions (Fig. 4C). Statistical comparisons were conducted pairwise between ΔFR values of different positions using independent-samples t-tests with FDR correction (Fig. 4D).

Given that each neuron has its own maximal ΔFR among the 13 positions, we focused on statistical comparisons between the maximum ΔFR and each other position for that neuron. This resulted in 13 t-tests per neuron, yielding a corresponding vector of p-values. Positions exhibiting significantly lower ΔFR than at the maximum, with p < 0.05, located before (i.e., at earlier time points than) the maximum position, defined the start boundary of the firing rate plateau as illustrated by the green line in Figures 5A–C. This indicates that ΔFR values at positions left (earlier) of this green line are significantly smaller than the maximum, whereas those between the green line and the maximum are not significantly different. Similarly, positions with significantly lower ΔFR than the maximum but occurring after (later than) the maximum position defined the end boundary of the plateau and were indicated by a blue line (Fig. 5A–C). Here, ΔFR values at positions right (later) of the blue line are significantly smaller than the maximum value, while those between the maximum and the blue line are not significantly different.

A “plateau period” was thus defined for each neuron as the continuous set of positions between these start (green line) and end (blue line) boundaries where ΔFR was statistically indistinguishable from the maximum ΔFR. To characterize the start and end effects further, the average ΔFR within the plateau period was computed. For the start effect, the ΔFR at the earliest position (e.g., 25 ms) was divided by this plateau average. For the end effect, the ΔFR at the latest position (e.g., 975 ms) was divided similarly. In cases where no significant difference was found at the farthest position (e.g., 975 ms, indicating no end effect), the plateau end position was set to that position by default. This quantitative measure captures the relative strength of neuronal modulation at boundary versus plateau positions. Statistical comparisons of firing rate ratios at the start time point were performed using independent t-tests within each brain region, whereas comparisons including both start and end time points across regions were assessed using ANOVA followed by appropriate post hoc tests.

Notably, we performed a one-tailed t-test against zero on this maximal ΔFR to determine whether the neuron exhibited a significant positive response. Neurons lacking a significant positive maximal ΔFR (p > 0.05) such as the examples shown in Supplementary Fig. 5 were excluded from the population analyses presented in Fig. 5. Consequently, in Fig. 5, data were recorded from 476 neurons in the A1, 495 neurons in the MGB, and 510 neurons in the IC.

#### LFP data analysis

Local field potentials (LFPs) were recorded simultaneously with single-unit activity using Neuropixels probes (initial sampling rate: 3 kHz). Raw voltage signals were bandpass filtered between 0.3 and 600 Hz, and notch filtered (second-order IIR) at 50/60 Hz and their harmonics to remove line noise. For each perturbation time, LFP traces were aligned to the perturbation onset, and a 0–100 ms post-change window was extracted for all subsequent analyses, consistent with the event-related windows used for EEG and ECoG.

Current Source Density (CSD) Computation. One-dimensional CSD profiles were computed from the second spatial derivative of the LFP (Pettersen, Devor et al. 2006, Szymanski, Garcia-Lazaro et al. 2009, Happel, Jeschke et al. 2010, El-Tabbal, Niekisch et al. 2021). The CSD at depth *z* was approximated by:

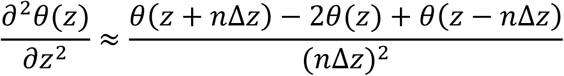

where θ is the field potential, Δz = 80 µm is the inter-channel spacing, and n = 2. A five-point Hamming spatial smoothing kernel (Happel, Jeschke et al. 2010) was applied to reduce high-frequency spatial noise.

Laminar boundaries were identified separately for each penetration. The earliest short-latency sink was designated as granular layer (Gr), with channels above and below assigned to supragranular (Sg) and infragranular (Ig) layers, respectively. CSD-derived sink patterns showed consistent agreement with previously published laminar profiles in rodent auditory cortex (Kaur, Rose et al. 2005, Sakata and Harris 2009, Szymanski, Garcia-Lazaro et al. 2009, Happel, Jeschke et al. 2010, Schaefer, Hechavarria et al. 2015, El-Tabbal, Niekisch et al. 2021). For population laminar plots (Fig. 6E), channels were resampled at 0.1-mm intervals within each penetration.

Time–frequency analysis. For each channel and perturbation time, the same 0–100 ms post-change window was used for time–frequency decomposition. Spectral power was computed using a Hann-tapered short-time Fourier transform (window length: 80 ms, 50% overlap) across 2–256 Hz (log-spaced). Change-related spectral modulation was quantified as:

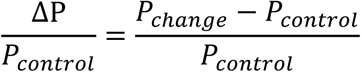

where P_control_ is the average power during no-change condition for the same channel.

Laminar aggregation for time–frequency responses (Fig. 7B). For each penetration, channels were grouped into Sg, Gr, and Ig based on CSD boundaries. Within each layer, the median ΔP/P_control_ across channels was computed, yielding one laminar profile per penetration per change time. These penetration-level medians were averaged across all penetrations (N = 17) to produce the laminar time–frequency maps.

Depth alignment across penetrations for band-limited responses (Fig. 7C). To compare spectral modulation across depths and penetrations, ΔP/P_control_ values were first averaged within canonical frequency bands: theta (4–8 Hz), alpha (8–14 Hz), beta (14–30 Hz), gamma (30–80 Hz), and high gamma (80–150 Hz). Depth alignment was then performed based on CSD-defined laminar boundaries: Sg channels were aligned by anchoring the deepest Sg channel across penetrations, Ig channels were aligned by anchoring the most superficial Ig channel, and Gr channels were center-aligned to the sink-defined centroid of the granular layer. Channels outside the overlapping depth range were excluded.

For each channel, change-related spectral power (ΔP/P_control_) was first averaged within a 0–100 ms post-change window and within predefined frequency bands. After laminar alignment, each depth-aligned channel contributed one band-limited ΔP/P_control_ value per perturbation position. These values were then averaged across penetrations to generate the depth-aligned laminar spectral profiles shown in Fig. 7C.

#### Computational Modeling

We implemented a mechanistic neural encoding model to simulate the time-course of sensitivity to local changes. The model consists of two sequential stages: a nonlinear encoding stage for mimicking spiking activity) and a temporal integration stage for mimicking LFP/population activity.

##### Stage 1: Neural Encoding and Saturation

The input stimulus was modeled as a step function (1000 ms duration) superimposed with a transient onset component (𝜏_𝑜𝑛𝑠𝑒𝑡_ = 30 𝑚𝑠) to mimic bottom-up saliency. Neural adaptation was modeled by a resource variable A(t) with a recovery time constant 𝜏_𝑎𝑑𝑎𝑝𝑡_ = 500 𝑚𝑠 , consistent with slow adaptation timescales in the auditory cortex (Ulanovsky, Las et al. 2004). The instantaneous firing rate R(t) was derived using the Naka-Rushton equation:

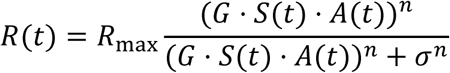

where 𝑛 = 3.0 controls the nonlinearity and 𝐺 represents baseline gain. The instantaneous encoding gain (proxy for spike-level sensitivity) was calculated as the derivative of the response with respect to the input (𝑑𝑅/𝑑𝑆). Parameters were tuned such that onset transients drove the response into the saturation plateau, minimizing the encoding gain.

##### Stage 2: Temporal Integration

To model cortical population dynamics, the instantaneous gain signal was convolved with a sliding integration window. The window width was set to 120 ms, consistent with the intrinsic timescales of sensory cortex (Murray, Bernacchia et al. 2014, Sabat, Gouyette et al. 2025) and psychophysical temporal summation windows. The integrated evidence (proxy for LFP/behavioral sensitivity) at time t was calculated as the moving average of the instantaneous gain over the interval [t, t + W].

The “end effect” emerged naturally as the moving window traversed the stimulus offset, incorporating post-stimulus silence (modeled with a 50-ms physiological decay constant) into the average.

## Acknowledgments

We are grateful to Prof. Nai Ding and Prof. Feixue Liang for their invaluable comments on an earlier version of the manuscript, as well as to Xiaokai Kou for their assistance with the experiments. This work was supported by STI2030-Major Projects (2022ZD0204800 and 2022ZD0204600) (to X.Y.); National Natural Science Foundation of China 32571216 and 32171044 (to X.Y.), 32100827 (to Yuying Zhai) and 32371050 (Yi Zhou).

---

**Supplementary Figure 1.**
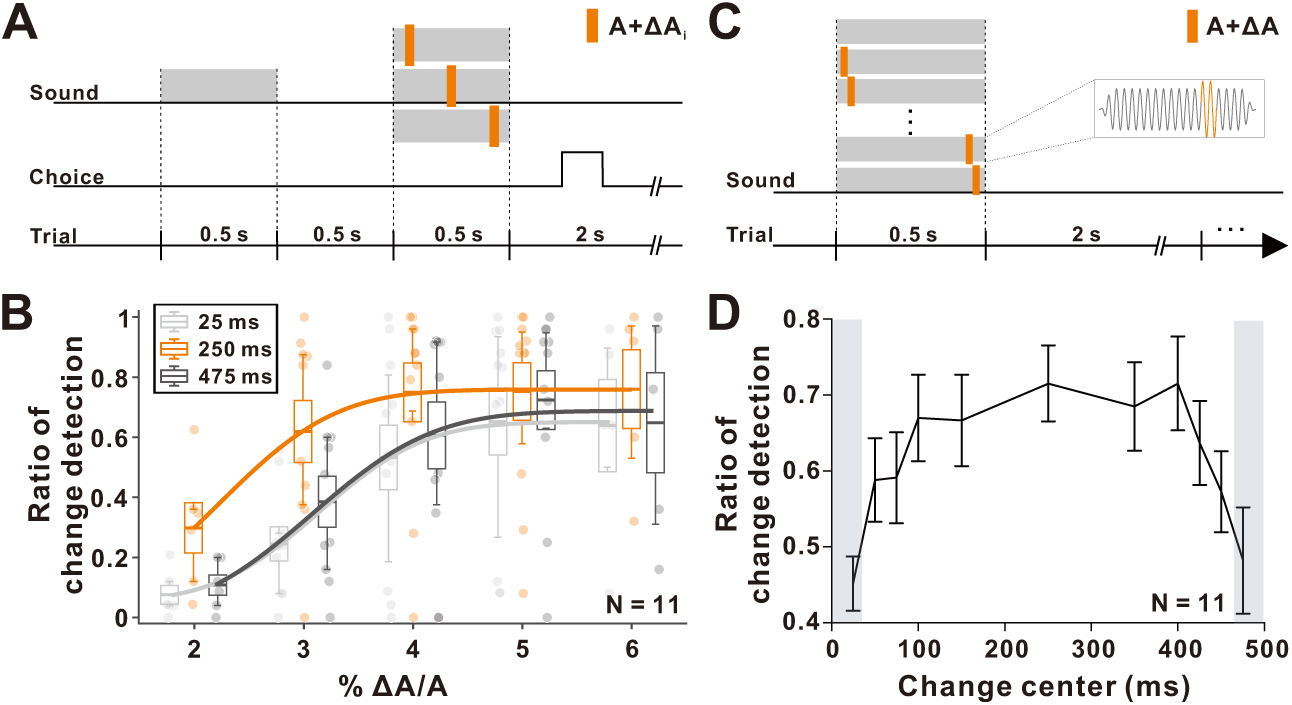
Psychological and behavioral results in the amplitude local change-detection task. **(A)** Schematic of the delayed match-to-sample task (Psychological Experiment, Amplitude Session 4). Participants compared two 500-ms, 1-kHz tones separated by a 500-ms silent interval. The second tone contained a brief amplitude increase (A + ΔA) occurring at 25 ms (light gray), 250 ms (orange), or 475 ms (dark gray) of its duration, with ΔA/A magnitude varied across trials (0%, 2%, 3%, 4%, 5%, 6%). Sixteen randomized stimuli were presented per session. Participants reported whether the two tones matched. **(B)** Change-detection performance for amplitude perturbations at 25 ms, 250 ms, and 475 ms. Transparent dots represent individual participants (N = 11); boxplots show the mean (central line), ±1 SEM (box edges), and 25th–75th percentiles (whiskers). Smooth curves show Gaussian fits to the pooled data. **(C)** Schematic of the amplitude change-detection task (Psychological Experiment, Amplitude Session 5). Participants heard a 500-ms, 1-kHz tone in which a brief local amplitude increase (A + ΔA) occurred at one of several temporal positions (vertical dashed lines: no-change, 25, 50, 75, 100, 150, 250, 350, 400, 425, 450, or 475 ms). Stimuli were randomly ordered across 12 conditions. Listeners indicated whether a change occurred via keypress. **(D)** Change-detection performance across temporal positions (N = 11). Error bars denote ±SEM. Light gray shaded rectangles mark temporal positions where detection performance was significantly lower than the position with maximal mean performance (one-way ANOVA across positions followed by post-hoc comparisons, p < 0.05), indicating reduced sensitivity near stimulus boundaries.

**Supplementary Figure 2.**
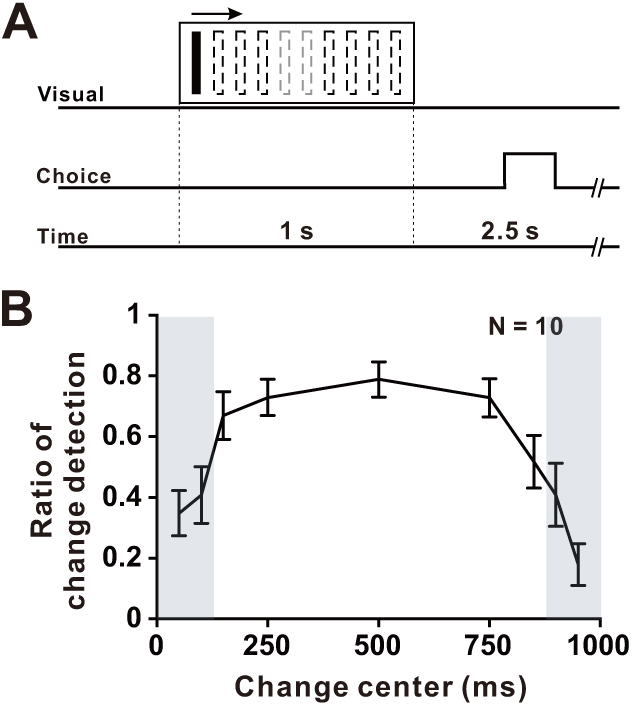
Psychological and behavior results in the visual local change-detection task. **(A)** Schematic of the visual change-detection paradigm (Psychological Experiment, Visual Session 6). Participants viewed a bar moving horizontally across the screen for 1000 ms. On each trial, a brief luminance increase could occur at one of several temporal positions (50, 100, 150, 250, 500, 750, 850, 900, or 950 ms), or no luminance change was presented (no-change condition). A total of 10 change conditions plus a no-change condition were presented in random order. After stimulus offset, participants indicated via keypress whether a luminance change had occurred. **(B)** Mean change-detection performance across temporal positions (N = 10). Error bars indicate ±SEM. Light gray shaded rectangles mark temporal positions where detection performance was significantly reduced relative to the position with maximal mean performance (one-way ANOVA across positions followed by post-hoc comparisons, p < 0.05), indicating reduced sensitivity near stimulus boundaries.

**Supplementary Figure 3.**
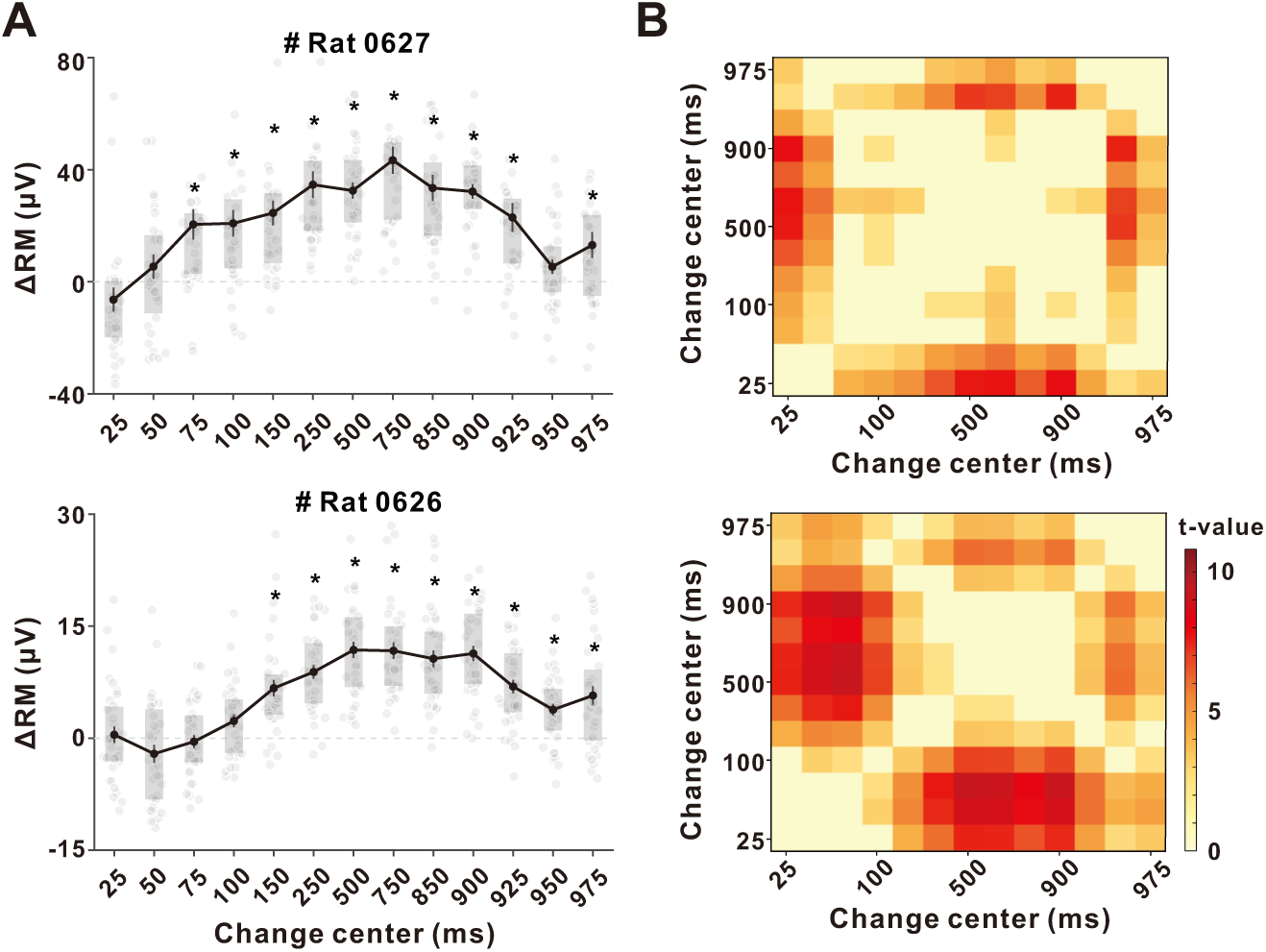
ECoG change responses during the local change-detection task in two additional rats. **(A)** ΔRM across perturbation positions for Rat 0627 (top) and Rat 0626 (bottom). For each rat, ΔRM values were averaged across the 16 channels exhibiting the strongest onset responses (see Methods). Gray dots represent ΔRM values from individual trials. Solid lines with filled circles denote the mean ΔRM across trials at each perturbation position, with vertical error bars indicating ±1 SEM. Shaded rectangles indicate the interquartile range (IQR; 25th–75th percentiles). The light gray dashed line denotes ΔRM = 0. Asterisks indicate perturbation positions where ΔRM was significantly greater than zero (one-tailed t-test, p < 0.05). **(B)** Pairwise t-value matrices comparing ΔRM across all perturbation positions for Rat 0627 (top) and Rat 0626 (bottom). Each matrix cell shows the t-value for the comparison between two perturbation times; nonsignificant entries (p ≥ 0.05) are masked. The color scale indicates the magnitude of the t-value, with darker colors reflecting stronger differences.

**Supplementary Figure 4.**
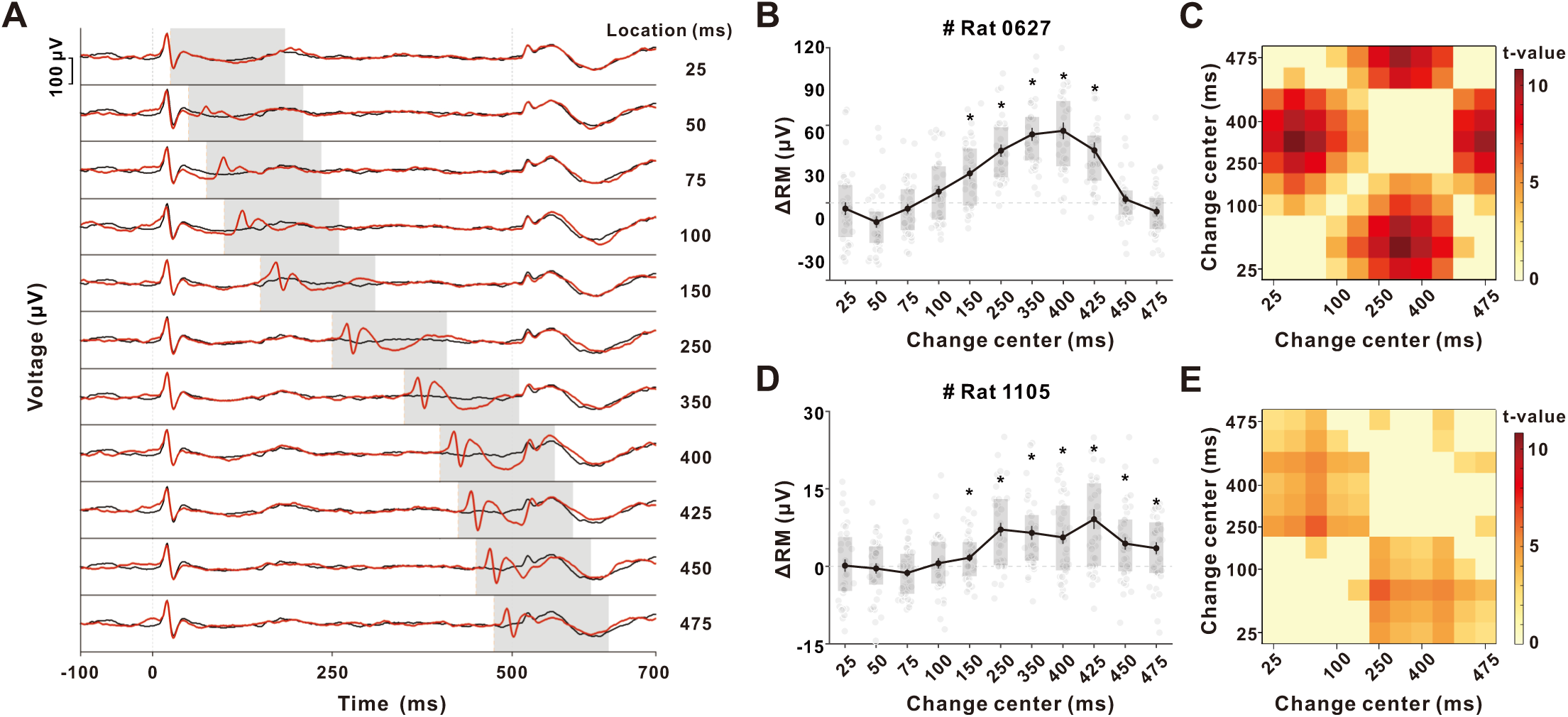
ECoG change responses during amplitude local change-detection task. **(A)** Example ECoG responses from channel #27 of Rat 0627 during the amplitude-change detection task (ECoG Experiment, Session 2). Black traces show responses in the control condition and red traces correspond to amplitude changes occurring at different perturbation positions. Orange dashed vertical lines indicate the tested perturbation times (25–475 ms) and gray dashed lines mark sound onset and offset. Shaded regions denote the analysis window (0–160 ms) used to compute RM, selected based on significant differences between the control and the 50% amplitude-change condition (permutation test, p < 0.05). **(B, D)** ΔRM across perturbation positions for Rat 0627 (B) and Rat 1105 (D). For each rat, ΔRM values were averaged across the 16 channels exhibiting the strongest onset responses (see Methods). Gray dots represent ΔRM values from individual trials. Solid lines with filled circles denote the mean ΔRM across trials at each perturbation position, with vertical error bars indicating ±SEM. Shaded rectangles indicate the IQR with 25th–75th percentiles. The light gray dashed line denotes ΔRM = 0. Asterisks indicate perturbation positions where ΔRM was significantly greater than zero (one-tailed t-test, p < 0.05). **(C, E)** Pairwise t-value matrices comparing ΔRM across all perturbation positions for Rat 0627 (C) and Rat 1105 (E). Each matrix cell reflects the t-value for the comparison between two perturbation times; nonsignificant entries (p ≥ 0.05) are masked. The color scale indicates t-value magnitude, with darker colors reflecting stronger differences.

**Supplementary Figure 5.**
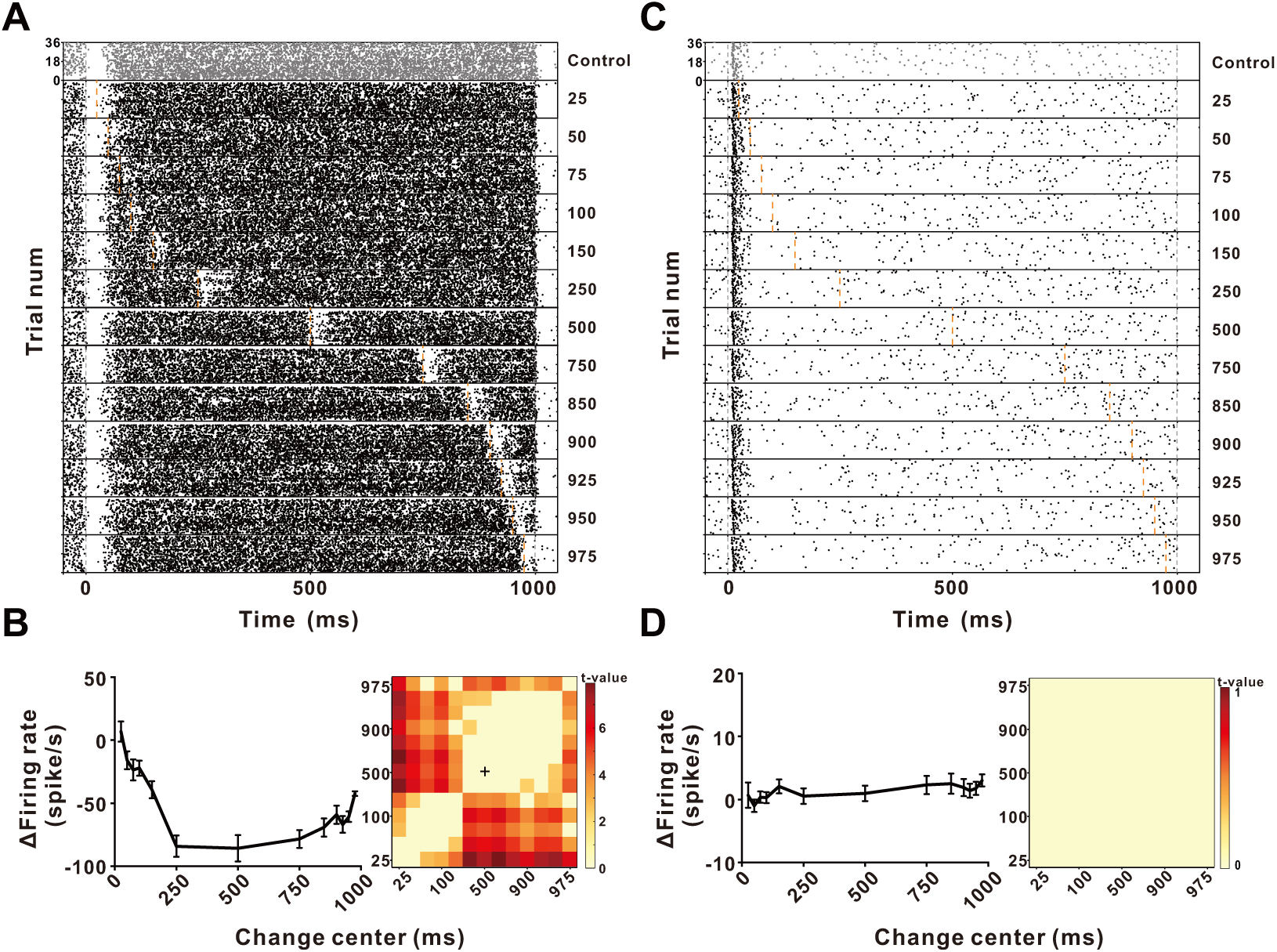
Example neuronal responses in the inferior colliculus (IC) to local changes. **(A)** Raster plot from an example IC neuron recorded during the local change-detection task (Extracellular Experiment). Orange dashed lines mark the perturbation times (25–975 ms) and gray vertical dashed lines indicate sound onset and offset. This neuron showed strong suppression to early changes followed by gradual recovery at later positions. **(B)** Left: Δ firing rate (ΔFR; change minus control, computed in a 0–100-ms window after perturbation) as a function of perturbation time. ΔFR exhibited pronounced suppression for early changes, reached a minimum around 500 ms, and then partially recovered toward later positions. Error bars indicate ±SEM. Right: Pairwise t-value matrix comparing ΔFR across positions (non-significant comparisons masked). The cross indicates the position of the minimum ΔFR. **(C–D)** A second IC neuron shown in the same format as panels (A–B). For this neuron, ΔFR remained near zero across all perturbation times, and the pairwise t-value matrix showed no significant differences, indicating an absence of systematic change-related modulation in this neuronal type.

**Supplementary Figure 6.**
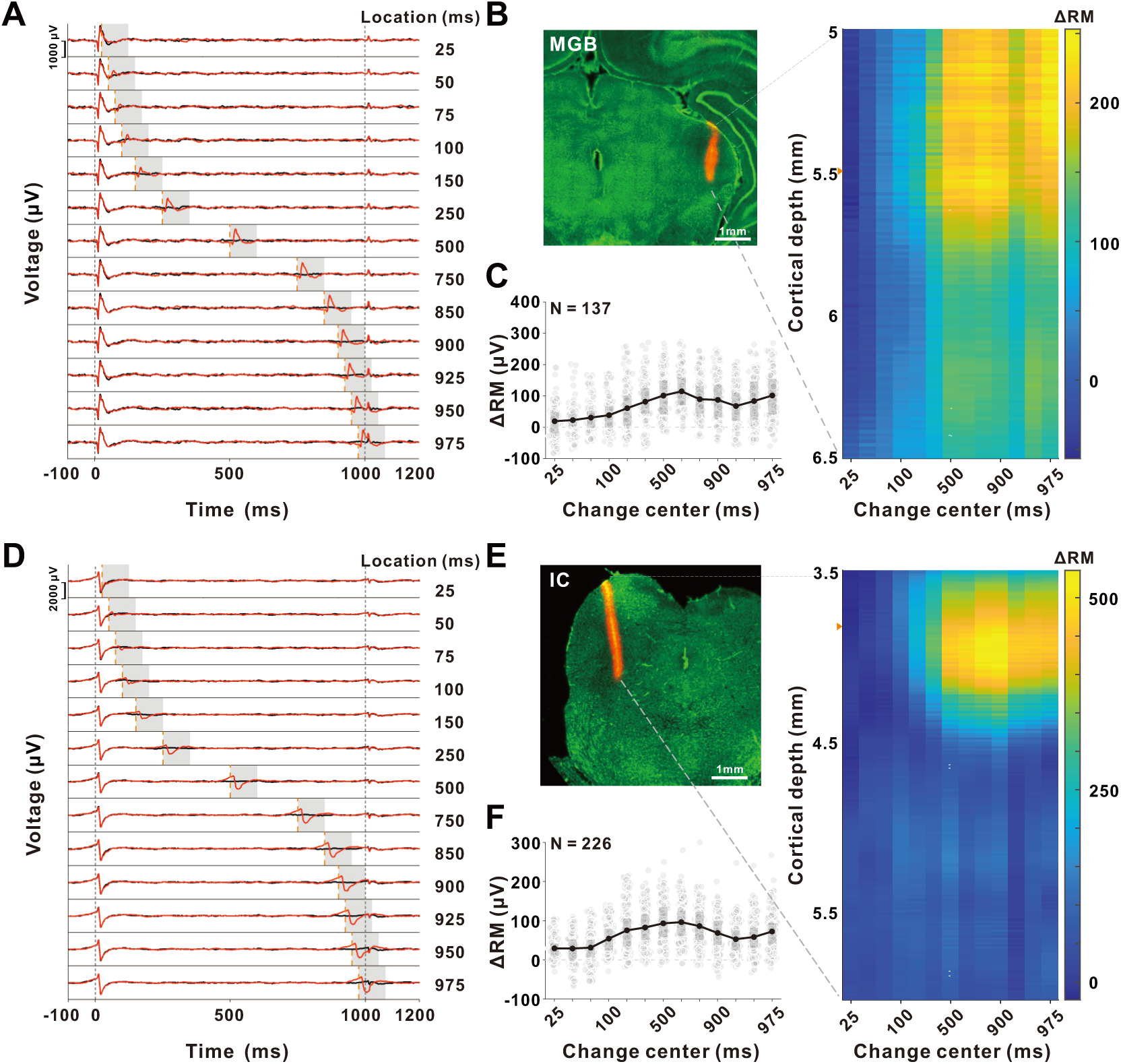
Laminar LFP responses to local change detection in MGB and IC. **(A)** Example LFP responses from a single MGB channel during the local change-detection task. Black traces show control responses; red traces show responses when a local change occurred at one of 13 temporal positions (25–975 ms). Gray shaded regions indicate the 0–100 ms analysis window used to compute RM. Gray vertical dashed lines indicate sound onset and offset, and orange vertical dashed lines mark the timing of the local change center. The example channel was selected from the depth indicated in panels (B). **(B)** Left: Histological verification of the MGB recording site. The orange track marks the probe trajectory. Right: Depth-resolved ΔRM map from a representative penetration within MGB. ΔRM is defined as RM_change_– RM_control_, computed within the 0–100 ms window. Color scale reflects ΔRM values across probe depth and change positions. **(C)** Population ΔRM across all MGB penetrations. Channels were sampled at 0.1-mm intervals along each probe (total N = 137 channels). Gray dots represent ΔRM values from individual channels. Solid lines connect the mean ΔRM across channels at each change position, with filled circles indicating the mean values. Shaded rectangles denote the IQR (25th–75th percentiles) across channels. Unlike A1, ΔRM does not decrease at late change positions, indicating the absence of end-related suppression in MGB. **(D–F)** Same analyses for IC. (D) Example IC channel responses, shown as in (A). (E) Left: Histological confirmation of the IC recording site and probe track. Right: Depth-resolved ΔRM map from a representative IC penetration. (F) Population ΔRM across IC channels sampled at 0.1-mm intervals (total N = 226 channels). symbols and shaded regions are defined as in (C). As in MGB, IC ΔRM remains elevated at late positions, indicating that robust end-related suppression is not present at the midbrain level.

## Notes

### Competing Interest Statement

The authors have declared no competing interest.

